# Human embryo-informed niche design enables defined bioprinting of undifferentiated hiPSCs

**DOI:** 10.64898/2026.11.12.765488

**Authors:** Gaeun Ju, Boyoung Kim, Jiyoon Kim, Mi Jeong Lee³, Jongjin Choi, Hee-Sick Youn, Seung-Woo Cho³, Sung-Hwan Moon, Soah Lee

**Author notes:** Correspondence to: Soah Lee.

## Abstract

Bioprinting undifferentiated human induced pluripotent stem cells (hiPSCs) enables post-printing expansion and lineage specification within geometrically defined constructs, yet the biological cues required to sustain pluripotent cells in a printable 3D microenvironment remain poorly defined. To establish a biologically grounded design framework, we analyzed single-cell transcriptomes from 106 human embryos to identify niche factors associated with pluripotency maintenance. Integrating ligand–receptor inference, pathway enrichment, and published evidence, we identified collagen IV (COL4A1), laminin (LAMA1), fibronectin (FN1), and pleiotrophin (PTN) as candidate niche factors. A minimal alginate–collagen IV–laminin bioink (ACL) supported colony formation, pluripotency, proliferation, and trilineage potential of bioprinted hiPSCs comparably to the Matrigel^TM^-containing bioink. Incorporating PTN (ACLP) further improved proliferation and survival under deprivation of pluripotency-promoting factors. This improvement was associated with PTPRZ1-dependent AKT signaling, as PTPRZ1 knockdown reduced p-AKT, pluripotency marker expression, and cell viability. Finally, the defined bioink supported printing of three embryo-inspired geometries—inner cell mass, bilaminar disc, and primitive streak. Collectively, this study establishes a single-cell embryo-informed strategy for rational bioink design and enables Matrigel-free bioprinting of undifferentiated hiPSCs.

## Introduction

Three-dimensional (3D) bioprinting enables the spatially controlled deposition of cells and biomaterials to generate tissue constructs with defined geometry and cellular organization^1,2^. Human induced pluripotent stem cells (hiPSCs) are particularly attractive for biofabrication because of their extensive self-renewal capacity and broad developmental potential^3,4^. Most hiPSC-based bioprinting strategies first expand hiPSCs and differentiate them into lineage-specific cell types, such as cardiomyocytes, neural cells, chondrocytes, osteogenic cells, or endothelial cells, before incorporating them into tissue-specific bioinks^5–10^. The composition and mechanical properties of these bioinks are generally tailored to support the survival, maturation, and function of the differentiated cell population after printing^11–14^. This strategy has enabled the fabrication of a wide range of tissue constructs, including cardiac, neural, cartilage, bone, and vascular tissues^7,9,15–19^.

Bioprinting hiPSCs in their undifferentiated state offers a distinct paradigm. Rather than assembling tissues from pre-specified cell populations, this approach preserves a proliferative and developmentally competent cell source within the printed construct, enabling post-printing expansion and subsequent lineage specification^20–22^. Previous studies have established that undifferentiated hiPSCs can survive bioprinting, expand within 3D constructs, retain pluripotency-associated properties, and remain competent for subsequent differentiation^23–30^. However, the bioinks used for these applications have largely been selected empirically on the basis of printability, cytocompatibility, or performance in specific differentiation contexts, rather than designed to recapitulate the biological niche that sustains pluripotent cells. Consequently, the extracellular and soluble cues required to maintain undifferentiated hiPSCs within a printable 3D microenvironment remain poorly defined^20^.

This limitation is particularly evident in the frequent use of Matrigel or Geltrex, mouse sarcoma-derived basement membrane extracts that effectively support hiPSC survival and proliferation through a complex mixture of extracellular matrix proteins and bioactive factors^31,32^. Their xenogeneic origin, undefined composition, and batch-to-batch variability limit reproducibility and mechanistic control^33^. Taken together, the field has demonstrated how to print undifferentiated hiPSCs, but lacks a biologically grounded framework for defining the niche that should support them after printing.

Here, we sought to define the niche components required to support undifferentiated hiPSCs after bioprinting and to incorporate these signals into a chemically defined bioink. We hypothesized that the early human embryo could provide a biological blueprint for identifying the extracellular matrix and paracrine cues that support pluripotency of hiPSCs. We first analyzed single-cell transcriptomes of human embryos, applying ligand–receptor inference and pathway analysis to identify key candidate niche factors. We then evaluated whether these factors preserve hiPSC survival, proliferation, pluripotency, and multilineage potential after bioprinting. We also demonstrated the feasibility of printing complex 3D geometries using the defined bioink. This embryo-informed strategy enabled the development of a Matrigel-free bioink that supports undifferentiated hiPSCs within printed constructs and provides a biologically grounded framework for rational bioink design.

## Results

### 3.1 Integrated human embryo scRNA-seq data identify candidate ligands associated with pluripotency maintenance

To identify microenvironmental factors associated with pluripotency maintenance during early human embryonic development, we analyzed three previously published human embryonic scRNA-seq datasets spanning 6–14 days post-fertilization (DPF)^34–36^ **(Figure 1a, S1a)**. The integrated dataset contained 6,832 high-quality cells and 46,432 expressed genes. UMAP visualization resolved six major embryonic cell populations: primitive endoderm (PrE), epiblast (EPI), inner cell mass (ICM), cytotrophoblasts (CTBs), syncytiotrophoblasts (STBs), and extravillous trophoblasts (EVTs) **(Figure 1b,c, S1b,c)**.

**Figure 1:**
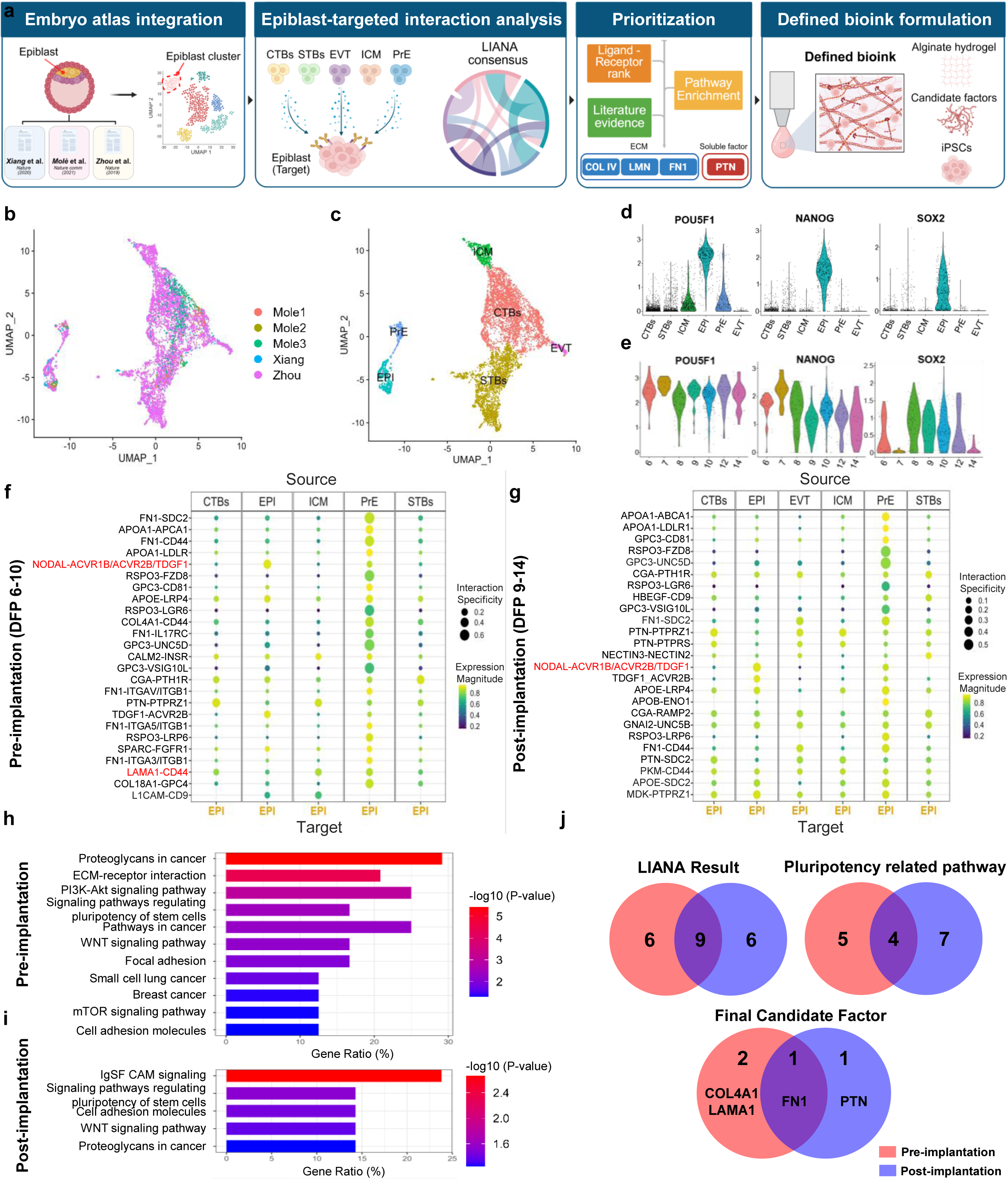
Integrative analysis of human embryonic scRNA-seq data identifies candidate niche factors for hiPSC bioink development. **a,** Schematic overview of the integration and analysis of three publicly available human embryonic single-cell RNA-sequencing (scRNA-seq) datasets spanning 6–14 days post-fertilization (DPF). **b,** UMAP visualization of the integrated dataset comprising 6,832 cells, colored by dataset of origin. **c,** Distribution of the six major embryonic cell populations: primitive endoderm (PrE), epiblast (EPI), inner cell mass (ICM), cytotrophoblasts (CTBs), syncytiotrophoblasts (STBs), and extravillous trophoblasts (EVTs). **d,** Expression of the pluripotency-associated markers *POU5F1/OCT4*, *NANOG*, and *SOX2* in the integrated dataset. **e,** Expression of pluripotency-associated markers in EPI cells across the developmental stages analyzed. **f,g,** Epiblast-targeted ligand–receptor interactions identified in pre-implantation and post-implantation developmental groups, showing conserved and stage-specific candidate ligands. **h,i,** KEGG pathway enrichment analysis of receptor genes corresponding to the selected epiblast-targeted ligand–receptor interactions. **j,** Stepwise prioritization of candidate ligands based on developmental conservation, pathway association, and previously reported functions in pluripotent stem-cell maintenance, survival, adhesion, or proliferation, resulting in the selection of COL4A1, LAMA1, FN1, and PTN for subsequent bioink development.

Because typical hiPSCs resemble the primed pluripotent state of the post-implantation epiblast, the epiblast population was selected for subsequent niche analysis^37,38^. Epiblast cells showed robust expression of the core pluripotency regulators POU5F1/OCT4, NANOG, and SOX2 (**Figure 1d; Figure S2**), with expression maintained across the developmental stages analyzed (**Figure 1e**).

To identify extracellular signals acting on the epiblast that could potentially support pluripotency, we analyzed epiblast-targeting ligand–receptor interactions. This analysis identified both conserved and stage-specific extracellular signals in the pre- and post-implantation groups **(Figure 1f,g)**. The top-ranked candidates included established pluripotency-associated factors such as NODAL and LAMA1^39–41^, supporting the validity of the analysis. The ranked candidates included both extracellular matrix-associated and soluble signaling molecules, with 9 ligands conserved across developmental stages (i.e., APOA1, APOE, CGA, FN1, GPC3, NODAL, PTN, RSPO3, TDGF1) and 6 ligands preferentially identified in each developmental group (Pre-implantation: CALM2, COL4A1, COL18A1, L1CAM, LAMA1, SPARC; Post-implantation: APOB, GNAI2, HBEGF, MDK, NECTIN3, PKM).

To distinguish ligand-receptor interactions potentially relevant to pluripotency maintenance from those associated with broader developmental processes, we next examined the signaling pathways associated with the corresponding epiblast receptors. KEGG pathway enrichment revealed pathways including “proteoglycans in cancer,” “ECM–receptor interaction,” “PI3K–Akt signaling,” “signaling pathways regulating pluripotency of stem cells,” and “Wnt signaling pathway” **(Figure 1h,i; Table S1)**. ECM-associated ligands, including FN1, COL4A1, and LAMA1, were linked to ECM–receptor and PI3K–Akt-related pathways, whereas soluble ligands, including NODAL, TDGF1, RSPO3, and PTN, were associated with pluripotency- and Wnt-related signaling pathways **(Table S1)**.

Based on these pathway associations, the candidate set was narrowed to 4 conserved (FN1, NODAL, RSPO3, TDGF1), 5 pre-implantation-specific (CALM2, COL4A1, COL18A1, LAMA1, SPARC), and 7 post-implantation-specific ligands (APOE, GNAI2, GPC3, MDK, NECTIN3, PKM, PTN) **(Figure 1j)**. Finally, we prioritized ligands with previously reported roles in the maintenance, survival, adhesion, or proliferation of human pluripotent stem cells, identifying COL4A1, LAMA1, FN1, and PTN as final candidates for subsequent bioink development^40,42–44^. Soluble factors that are already supplied by defined pluripotent stem cell culture media, including FGF2, TGF-β1, and NODAL, were excluded from further consideration^45^. Receptors corresponding to these candidate ligands, including FGFR1, PTPRZ1, ITGB1, ITGB5, and FZD5, were expressed in the epiblast population **(Figure S3a)**. qRT-PCR and RT-PCR further confirmed their endogenous expression in hiPSCs **(Figure S3b,c)**.

### 3.2 Alginate-based collagen IV and laminin bioink (ACL) supports pluripotency of bioprinted hiPSCs

To experimentally validate whether the bioinformatically identified candidate niche factors could promote hiPSC survival and pluripotency after bioprinting, we first focused on collagen IV and laminin, two extracellular matrix components identified in our analysis and major constituents of Matrigel^32,46^ **(Figure S4).** We used alginate hydrogel as the structural backbone of the bioink because of its biocompatibility, mild ionic crosslinking, and structural stability^47,48^. To evaluate whether a defined alginate-based bioink containing collagen IV and laminin could replace complex Matrigel, we compared alginate alone, alginate + collagen IV (AC), alginate + laminin (AL), alginate + collagen IV + laminin (ACL), and Matrigel-containing conditions over 4 days of post-printing culture in the droplet format (**Figure 2a**). Because each printed droplet represents the smallest structural unit, or voxel, of a bioprinted construct, we evaluated hiPSC proliferation and pluripotency within individual voxels.

**Figure 2:**
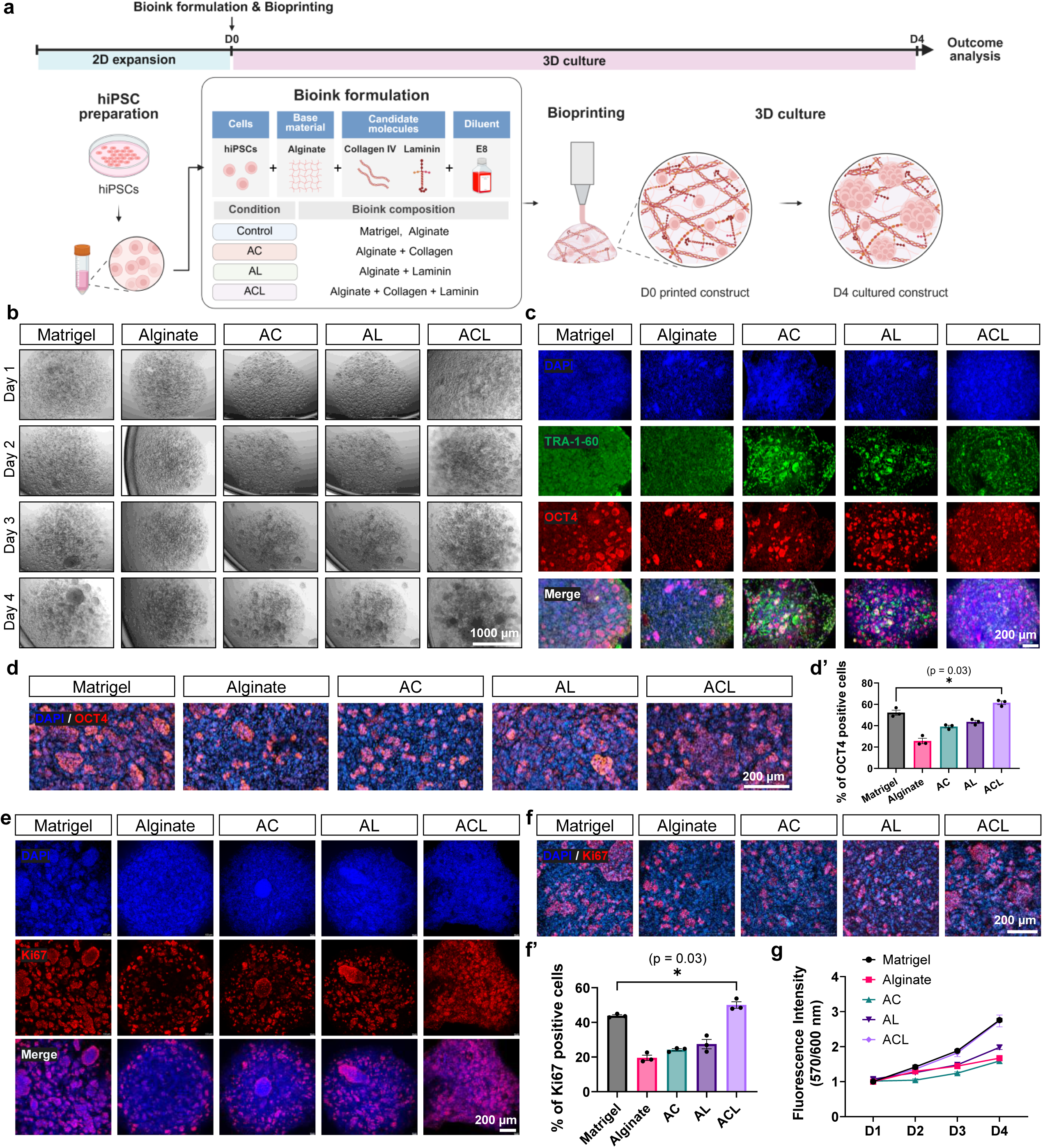
Alginate-based collagen IV and laminin bioink supports pluripotency-associated properties of bioprinted hiPSCs. **a,** Schematic of the experimental design for evaluating alginate-based bioink formulations containing collagen IV and/or laminin. hiPSCs were bioprinted in alginate alone, alginate with collagen IV (AC), alginate with laminin (AL), alginate with collagen IV and laminin (ACL), or a Matrigel-containing formulation and cultured for 4 days. **b,** Representative bright-field images of printed hiPSC-containing droplets during culture. **c,d,** Representative immunofluorescence images of OCT4 and TRA-1-60 expression in bioprinted hiPSCs on day 4. **d**′, Quantification of OCT4-positive cells. **e,f,** Representative immunofluorescence images of Ki67 expression in bioprinted hiPSCs on day 4. **f**′, Quantification of Ki67-positive cells. **g,** Longitudinal metabolic activity of bioprinted hiPSCs assessed using the PrestoBlue assay.

Bright-field imaging showed that the printed droplets retained their shape in all conditions throughout the culture period, with cells distributed across the droplet volume. Over time, a subset of cells formed small aggregates (**Figure 2b**). Immunofluorescence staining on day 4 showed clear expression of the pluripotency markers OCT4 and TRA-1-60 in all conditions, although to varying extents (**Figure 2c,d**). The proportion of OCT4-positive cells was significantly higher in the ACL condition (64.1 ± 5.1%) than in all other conditions, including the Matrigel positive control (57.7 ± 2.8%, p = 0.03 vs. Matrigel; Figure 2d’). Immunofluorescence staining for the proliferation marker Ki67 showed its robust expression in hiPSCs cultured within the ACL bioink (**Figure 2e,f**). Quantification showed that the percentage of Ki67-positive cells in the ACL condition was significantly higher than that in the Matrigel condition (50.0 ± 1.9% vs. 43.8 ± 0.6%, p = 0.03; **Figure 2f’**). Longitudinal analysis of metabolic activity using the PrestoBlue assay showed a progressive increase in metabolic activity in the ACL condition, comparable to the Matrigel condition and higher than that observed in the single-ECM conditions (**Figure 2g**). Together, the ACL bioink supported hiPSC aggregation, proliferation, metabolic activity, and pluripotency marker expression within alginate-based 3D-printed constructs at levels comparable to or greater than those in the Matrigel condition.

To characterize the Matrigel benchmark used in these comparisons, we performed proteomic profiling of three independent production lots. A relatively conserved matrisome composition was observed, with 502 proteins shared across all three lots **(Figure S4a)** and similar core matrisome and matrisome-associated protein profiles **(Figure S4b–e)**. Laminin-family proteins, including LAMA1, LAMB1, and LAMC1, were consistently abundant within the glycoprotein fraction, and type IV collagen, particularly COL4A2 together with COL4A1, was consistently detected within the collagen fraction **(Figure S4c, f–g),** confirming that the two components of the ACL bioink were abundant constituents of Matrigel. Notably, non-matrisome proteins accounted for approximately 17–19% of the detected protein abundance and included several intracellular proteins **(Figure S4b, j–k)**, underscoring the undefined nature of this benchmark matrix. Consistent with the compositional similarity across lots, no significant differences in the measured biological responses were detected among the three Matrigel lots **(Figure S5)**.

### 3.3 Supplementation of ACL bioink with fibronectin and pleiotrophin preserves pluripotency of bioprinted hiPSCs

After confirming that the ACL maintained pluripotency comparably to Matrigel, we further refined the ACL formulation by incorporating two additional candidate niche factors identified from our analysis: fibronectin (FN1), an ECM molecule, and pleiotrophin (PTN), a soluble factor. Dose–response analysis identified 100 μg/ml FN1 and 50 ng/ml PTN as the concentrations producing the highest Ki67-positive cell fractions among the tested doses **(Figure S6)**. These concentrations were therefore used for subsequent formulation studies.

The tested formulations included ACL, ACL supplemented with FN1 (ACLF), PTN (ACLP), or both FN1 and PTN (ACLFP). Bright-field microscopy showed stable hiPSC aggregate formation across all formulations (**Figure 3a**). To examine whether supplementation with FN1 and PTN influenced the pluripotency of bioprinted hiPSCs, changes in OCT4 and TRA-1-60 expression were analyzed. Both low- and high-magnification images showed high OCT4 expression across all conditions (Figure 3b–c’). The percentage of OCT4-positive cells was slightly higher in ACLP than in ACL (69.9 ± 2.5% vs. 64.0 ± 5.1%), although this difference was not statistically significant (p = 0.5337). To further examine the effect of FN1 and PTN supplementation, we assessed hiPSC proliferative activity by Ki67 immunofluorescence staining (Figure 3d–e’). Formulations containing FN1, with or without PTN (ACLF and ACLFP), showed proliferation levels comparable to ACL. By contrast, the ACLP formulation significantly increased the percentage of Ki67-positive cells relative to ACL (60.3 ± 2.7% vs. 49.3 ± 2.2%; p = 0.0333; **Figure 3e’**). PrestoBlue-based longitudinal analysis demonstrated a continuous increase in metabolic activity across all formulations (**Figure 3f**). By day 4, ACL, ACLF, and ACLP showed comparable metabolic activity, with no significant difference between ACL and ACLP (p = 0.7489), whereas ACLFP showed a modest decrease in fluorescence intensity.

**Figure 3:**
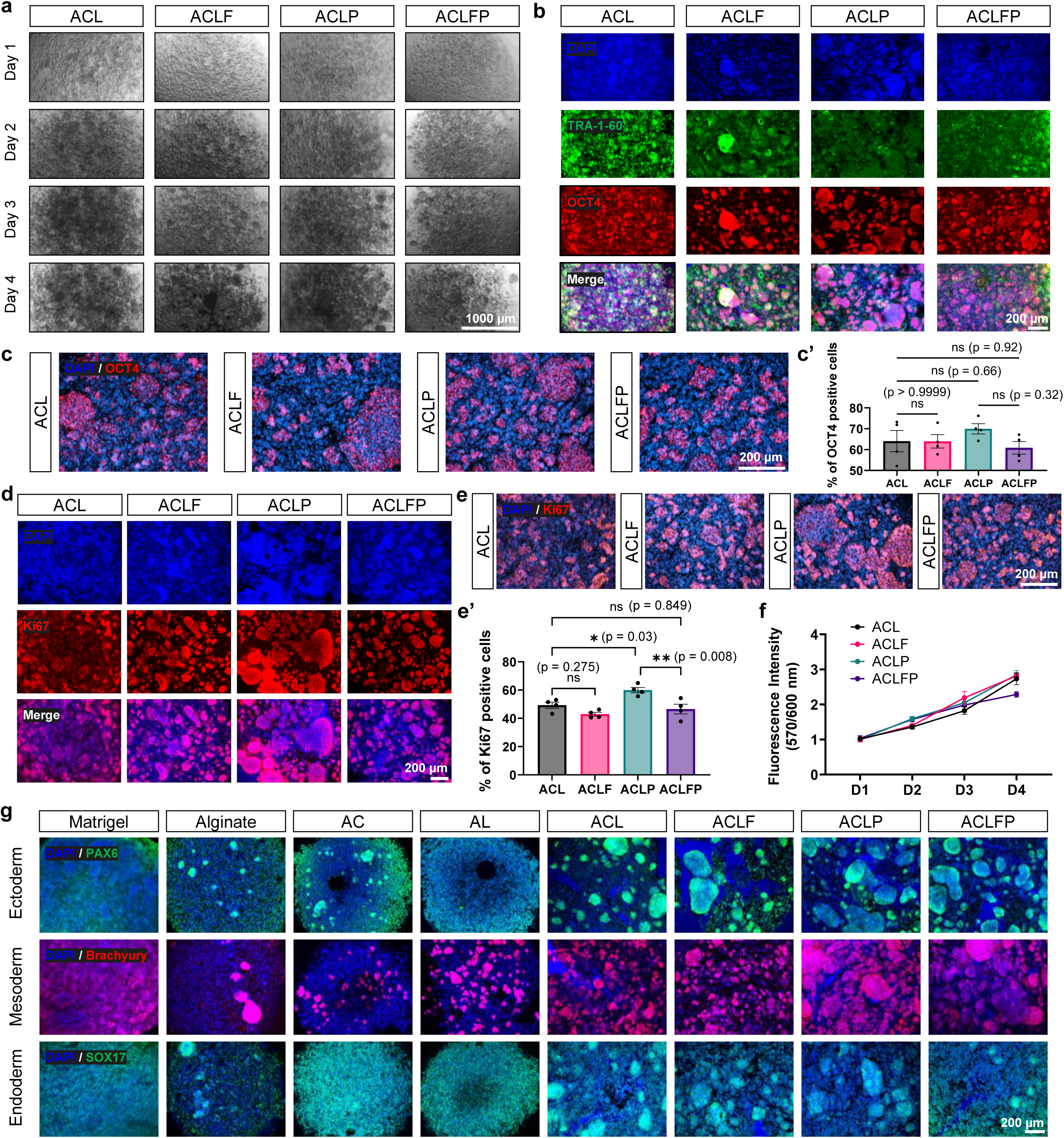
Pleiotrophin supplementation enhances proliferative activity in ACL-based bioinks while preserving pluripotency-associated properties of bioprinted hiPSCs. **a,** Representative bright-field images of bioprinted hiPSCs cultured in ACL alone or in ACL supplemented with fibronectin (ACLF), pleiotrophin (ACLP), or both (ACLFP). **b,c,** Representative immunofluorescence images of OCT4 and TRA-1-60 expression in the indicated bioink formulations on day 4. **c**′, Quantification of OCT4-positive cells. **d,e,** Representative immunofluorescence images of Ki67 expression in the indicated formulations on day 4. **e**′, Quantification of Ki67-positive cells. **f,** Longitudinal metabolic activity measured using the PrestoBlue assay. **g,** Immunofluorescence analysis following directed differentiation toward the three germ layers, showing expression of PAX6, Brachyury, and SOX17 as markers of ectoderm, mesoderm, and endoderm, respectively.

Finally, we performed directed differentiation toward the three germ layers. Immunofluorescence staining confirmed the expression of the ectoderm marker (PAX6), mesoderm marker (Brachyury), and endoderm marker (SOX17) across all conditions, demonstrating that the hiPSCs retained multilineage differentiation potential (**Figure 3g**). Taken together, the ACL-based formulations supported hiPSC aggregation, pluripotency-associated marker expression, proliferation, metabolic activity, and multilineage differentiation potential, while PTN supplementation selectively enhanced proliferative activity.

Because the ACLP formulation supported proliferation, metabolic activity, and pluripotency-associated marker expression, we selected it as the lead candidate and assessed pluripotency *in vivo*. hiPSCs were harvested from droplet-printed ACLP constructs on day 4 and injected into the right testis of immunodeficient mice. At 16 weeks post-injection, the bioprinted hiPSCs formed teratomas containing derivatives of all three germ layers, including neural rosettes, cartilage, and gut epithelium **(Figure S7)**, indicating that hiPSCs retained functional pluripotency after bioprinting in ACLP bioink.

### 3.4 PTN contributes to preserving hiPSC pluripotency via the PTPRZ1–AKT signaling axis

In the preceding experiments, E8 medium was used as the bioink diluent. Although PTN supplementation increased hiPSC proliferation, its effect on pluripotency marker expression was limited, potentially because E8 contains pluripotency-supporting factors such as FGF2 and TGF-β^45^. To more clearly evaluate the contribution of PTN, ACL and ACLP bioinks were prepared using basal DMEM/F12 lacking these exogenous growth factors, while the printed constructs were subsequently cultured in E8 medium for 4 days **(Figure 4a).**

**Figure 4:**
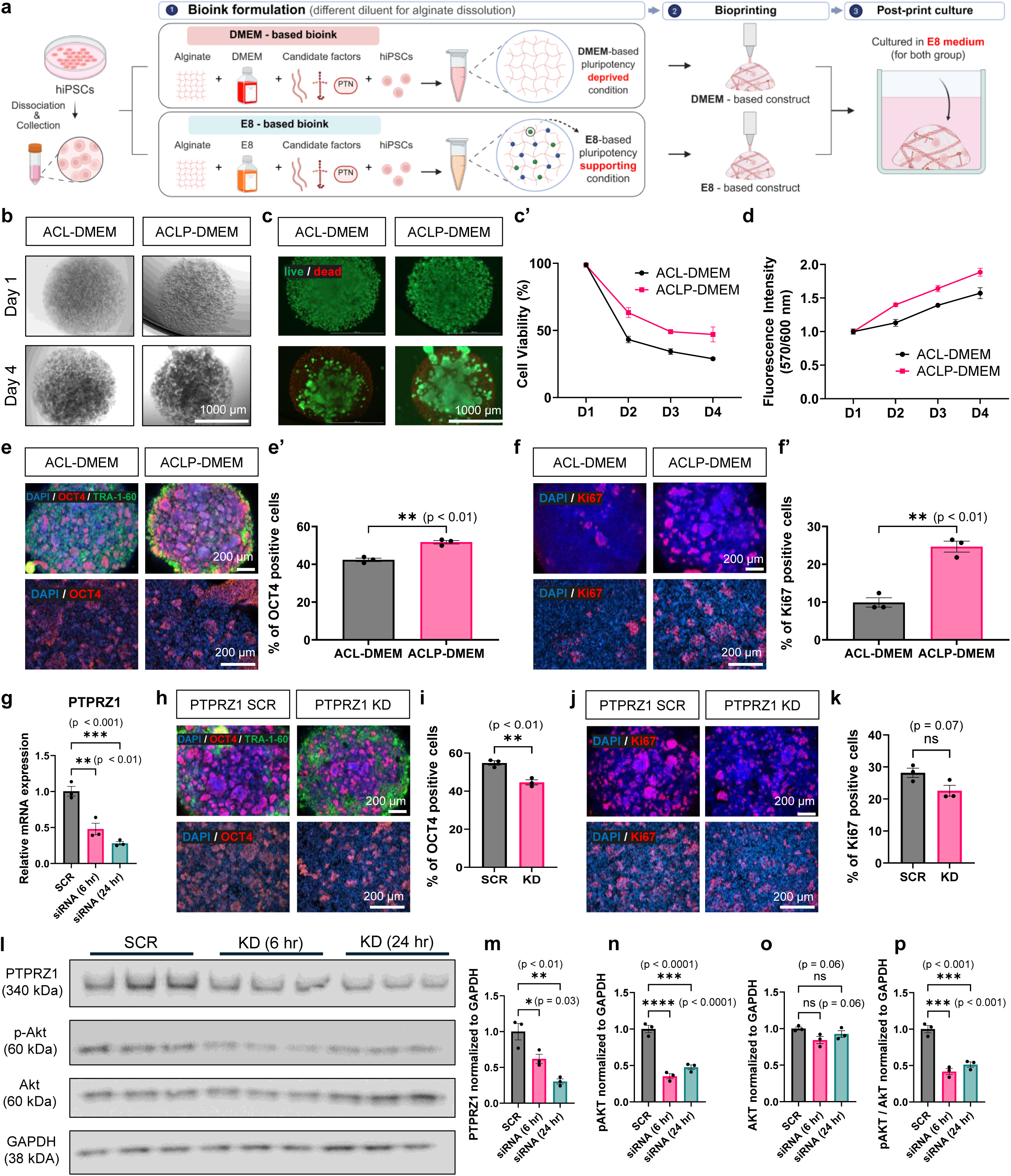
PTN contributes to the maintenance of pluripotency-associated properties through PTPRZ1–AKT signaling in bioprinted hiPSCs. **a,** Schematic of the experimental design used to assess the contribution of PTN under different bioink diluent conditions. ACL and ACLP bioinks were prepared using E8 medium or basal DMEM/F12 lacking exogenous pluripotency-supporting growth factors, followed by culture of the printed constructs in E8 medium. **b,** Representative bright-field images of hiPSC aggregates in DMEM-ACL and DMEM-ACLP conditions. **c,** Representative live/dead fluorescence images on day 4. **c**′, Quantification of cell viability. **d,** Longitudinal metabolic activity assessed using the PrestoBlue assay. **e,** Representative immunofluorescence images of OCT4 expression in DMEM-ACL and DMEM-ACLP constructs on day 4. **e**′, Quantification of OCT4-positive cells. **f,** Representative immunofluorescence images of Ki67 expression. **f**′, Quantification of Ki67-positive cells. **g,** Relative *PTPRZ1* mRNA expression following transfection with scrambled control or PTPRZ1-targeting siRNA at the indicated time points. **h,** Representative OCT4 immunofluorescence images following PTPRZ1 knockdown. **i,** Quantification of OCT4-positive cells. **j,** Representative Ki67 immunofluorescence images following PTPRZ1 knockdown. **k,** Quantification of Ki67-positive cells. **l,m,** Western blot analysis and densitometric quantification of PTPRZ1 protein following PTPRZ1 knockdown. **n–p,** Western blot analysis and densitometric quantification of phosphorylated AKT (p-AKT) relative to total AKT following PTPRZ1 knockdown at the indicated time points.

When DMEM/F12 was used as the bioink diluent, ACLP promoted hiPSC aggregate formation and cell viability compared with ACL **(Figure 4b,c).** Cell viability on day 4 was 28.9 ± 2.0% in DMEM-ACL and 47.0 ± 5.5% in DMEM-ACLP (**Figure 4c’**). Consistently, the PrestoBlue assay showed higher metabolic activity in DMEM-ACLP than in DMEM-ACL throughout the culture period (**Figure 4d**). Immunofluorescence staining showed that the percentage of OCT4-positive cells was significantly higher in DMEM-ACLP than in DMEM-ACL (51.8 ± 0.9% vs. 42.4 ± 0.9%, p = 0.0018; **Figure 4e,e’**). Ki67-positive cells were 2.49-fold higher in DMEM-ACLP than in DMEM-ACL (p = 0.0015; **Figure 4f,f’**). These results indicate that PTN supplementation partially preserved viability, proliferative activity, and pluripotency-associated marker expression even when pluripotency-supporting factors were omitted from the bioink diluent.

Protein tyrosine phosphatase receptor type Z1 (PTPRZ1) is a well-established receptor for PTN^49^ and was predicted to be the corresponding receptor on the epiblast in our ligand–receptor analysis (**Figure 1g**). To determine whether the effects of PTN were mediated through PTPRZ1, we knocked down PTPRZ1 in hiPSCs using PTPRZ1-specific siRNA. qRT-PCR showed that PTPRZ1 mRNA decreased in a time-dependent manner relative to the scrambled control, corresponding to approximately 52% knockdown at 6 h (p < 0.01) and 72% at 24 h (p < 0.001) (**Figure 4g**). Bright-field and live/dead imaging showed less compact aggregates and a greater dead-cell signal after PTPRZ1 knockdown (**Figure S8a,b**). Cell viability declined over the 4-day culture in both groups but fell more steeply after knockdown, reaching 27.7 ± 2.8% on day 4 compared with 45.3 ± 1.3% in the scrambled control (p = 0.004; **Figure S8c**). Metabolic activity increased over the culture period in both groups, but the increase was smaller after knockdown, with the two curves diverging from day 3 onward (**Figure S8d**). With regard to supporting pluripotency, PTPRZ1 knockdown decreased the OCT4-positive fraction from 54.8 ± 1.2% to 44.6 ± 1.3% (p = 0.0044; **Figure 4h,i**). The Ki67-positive fraction showed a similar decreasing trend, from 28.2 ± 1.5% to 22.6 ± 1.7%, although this difference did not reach statistical significance (p = 0.0671; **Figure 4j,k**). Together, these results implicate the involvement of PTPRZ1 in the PTN-associated maintenance of hiPSC viability and pluripotency marker expression after bioprinting.

Previous studies have shown that PTN promotes hESC survival and clonal growth via PTPRZ1, partly through AKT^43^, and AKT activation is sufficient to maintain pluripotency in mouse and primate ESCs^50^. We therefore examined whether PTN-PTPRZ1 signaling leads to AKT activation in bioprinted hiPSCs. PTPRZ1 knockdown significantly reduced PTPRZ1 protein levels to 0.62 ± 0.06 at 6 h and 0.30 ± 0.04 at 24 h relative to the scrambled control (p = 0.0257 and p = 0.0015, respectively; **Figure 4l,m**). Consistently, the p-AKT/total AKT ratio was significantly reduced to 0.42-fold at 6 h and to 0.51-fold at 24 h relative to the scrambled control (p = 0.0001 and p = 0.0004, respectively; **Figure 4n–p**).

To further investigate signaling downstream of AKT, we analyzed β-catenin/Wnt activity. β-catenin/Wnt signaling is implicated in stem cell self-renewal and pluripotency-associated cellular states^51,52^ and was also identified in our KEGG pathway enrichment analysis of epiblast-expressed receptor genes (**Figures 1h,i; Table S1**). Consistent with the western blot data (**Figure 4l,m**), immunostaining showed reduced PTPRZ1 protein after knockdown (**Figure S9a**). In the same construct, PTPRZ1 knockdown significantly decreased the nuclear-to-cytoplasmic ratio of β-catenin compared with the scrambled control group (p = 0.002; **Figure S9b,c**). In line with reduced nuclear β-catenin available for transcriptional co-activation, qRT-PCR showed decreased expression of β-catenin/Wnt downstream target genes, including AXIN2, LEF1, and CCND2 (p < 0.02; **Figure S9d**). Based on the established regulation of β-catenin signaling by AKT through GSK3β-dependent mechanisms^53–55^, these findings link reduced AKT activation to decreased β-catenin/Wnt activity after PTPRZ1 knockdown (**Figure S9e,f**), implicating PTN-PTPRZ1 signaling in the maintenance of pluripotency and proliferative activity in bioprinted hiPSCs.

### 3.5 The defined bioink enables fabrication of structurally stable, embryo-inspired hiPSC constructs that retain pluripotency

Having established that the defined ACLP bioink is printable and supports hiPSC viability, proliferation, and pluripotency at the single-voxel (droplet) scale, we next asked whether it could build more complex geometries. Three structures inspired by morphological features of early human development—inner cell mass (ICM), bilaminar disc, and primitive streak—were designed using computer-aided design (CAD) **(Figure 5a).** We first assessed dimensional fidelity under acellular conditions, defined as the ratio of each measured construct dimension to the corresponding CAD value. All three constructs reproduced the overall geometry of the original designs, with most dimensional fidelity values remaining close to the reference value of 1 **(Figure 5b).** Significant deviations were observed in the wall thickness of the ICM and bilaminar disc constructs (p = 0.002 for both) and in the height of the primitive streak construct (p = 0.027), whereas the remaining dimensions did not differ significantly from the corresponding CAD values. These results indicate that the defined bioink reproduced the overall geometry of the three embryo-inspired designs, although not all dimensions matched the design values.

**Figure 5:**
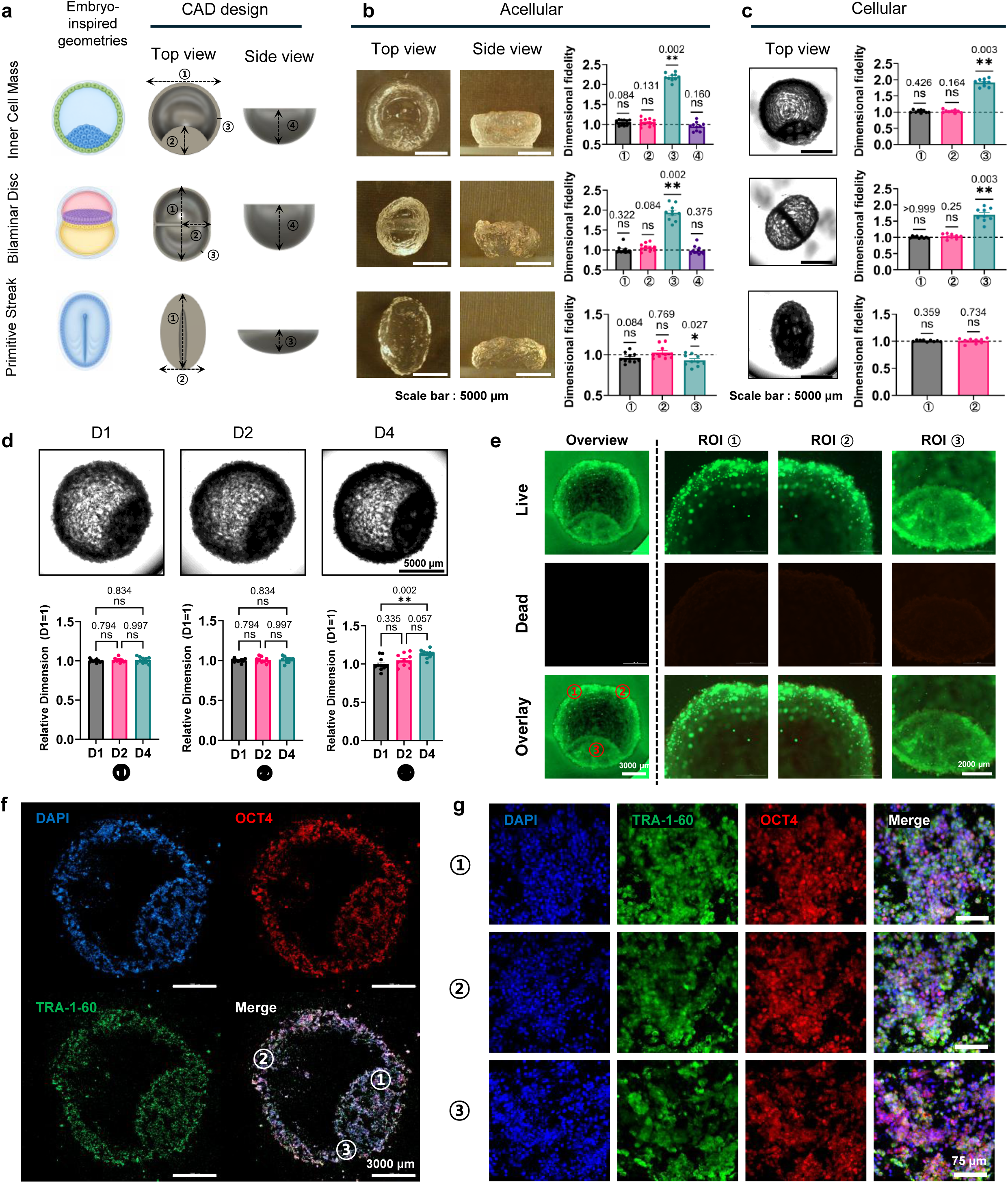
Defined ACLP bioink enables fabrication of embryo-inspired hiPSC constructs with short-term structural stability and maintenance of pluripotency-associated marker expression. **a,** Computer-aided design (CAD) models of three embryo-inspired structures representing the inner cell mass (ICM), bilaminar disc, and primitive streak, shown in top and side views. **b,** Representative images and dimensional fidelity analysis of acellular ACLP constructs fabricated using the corresponding CAD designs. Dimensional fidelity was calculated as the ratio of each measured construct dimension to its CAD-defined value. **c,** Representative top-view images and dimensional fidelity analysis of the corresponding hiPSC-containing constructs. **d,** Structural stability of printed ICM constructs during culture on days 1, 2, and 4. Individual dimensions were normalized to their respective day-1 values. **e,** Live/dead staining of day-4 ICM constructs showing the whole construct and representative regions of interest. **f,** Immunofluorescence staining of cryosectioned day-4 ICM constructs for OCT4 and TRA-1-60. **g,** Higher-magnification images of three representative regions within the ICM construct showing OCT4- and TRA-1-60-positive cells.

We next incorporated hiPSCs into the bioink and fabricated the same constructs under cellular printing conditions. Because the cellular constructs deformed when removed from the culture medium, side-view imaging was not feasible. Dimensional fidelity was therefore assessed only from top-view measurements of the immersed constructs (**Figure 5c**). Most dimensions were comparable to their CAD-defined values. Consistent with the acellular data, the wall thickness of the ICM and bilaminar disc constructs again exceeded the CAD value (p = 0.003 for both). No significant differences were observed in the remaining dimensions (all p ≥ 0.25). Thus, the bioink supported fabrication of spatially distinct geometries in the presence of encapsulated hiPSCs.

To assess structural stability during culture, the bioprinted ICM constructs were monitored on days 1, 2, and 4 (**Figure 5d**). Individual dimensions were normalized to their respective day-1 values. Dimensions ⍰and ⍰remained relatively stable, whereas dimension showed a modest but significant increase by day 4 (p < 0.01). Despite this localized change, the overall top-view morphology was maintained through day 4, indicating short-term structural stability.

We next assessed cell survival within the printed structures by performing live/dead staining of day-4 ICM constructs. Imaging of the entire construct and three spatially distinct regions showed predominantly live-cell fluorescence with limited dead-cell signal **(Figure 5e),** indicating that hiPSCs remained viable throughout the mm-scale constructs through day 4.

Finally, to determine whether the printed hiPSCs retained pluripotency-associated marker expression within the printed constructs, day-4 ICM constructs were sectioned and immunostained for OCT4 and TRA-1-60 **(Figure 5f).** Both markers were detected throughout the sections, and higher-magnification imaging of three representative regions revealed OCT4- and TRA-1-60-positive cells at each location **(Figure 5g).** Collectively, these findings show that the defined ACLP bioink enables fabrication and short-term maintenance of embryo-inspired 3D structures while supporting cell viability and pluripotency marker expression.

## Discussion

We identified a minimal, chemically defined combination of collagen IV, laminin, and PTN for maintaining undifferentiated hiPSCs after bioprinting at levels comparable to Matrigel (**Figures 1–3**). Our findings support the use of the early human embryo as a biological reference for prioritizing extracellular and soluble cues that may contribute to maintenance of the pluripotent state (**Figure 1**).

Defined alternatives to Matrigel have been extensively developed for pluripotent stem cell culture. In two-dimensional systems, recombinant vitronectin, laminin-511/521, laminin E8 fragments, and synthetic peptide- or polymer-based substrates have supported long-term self-renewal under chemically defined conditions^41,56–59^. In three-dimensional culture, hyaluronic acid-, PEG-, and other synthetic hydrogel systems, including matrices functionalized with integrin-binding peptides, have further demonstrated that pluripotency can be maintained within defined 3D environments while allowing control over adhesive and mechanical properties^60–62^. Related studies in undifferentiated PSC bioprinting have shown that pluripotent cells can survive printing, expand within 3D constructs, retain pluripotency-associated properties, and subsequently differentiate using bioinks based on alginate, chitosan, agarose, cellulose derivatives, or Matrigel-containing formulations^23,25,30,63^. Collectively, these studies established the feasibility of defined PSC culture and undifferentiated PSC bioprinting, while our study extends this progression by using developmental information to guide selection of biochemical cues for a printable pluripotent cell microenvironment.

This design strategy identified laminin, collagen IV, and PTN as a minimal combination supporting undifferentiated hiPSCs after bioprinting. Importantly, their effects were not simply additive. FN1 provided no additional benefit to the ACL matrix and reduced both viability and pluripotency when added to ACLP (**Figure 3**). Thus, identification of a factor as part of a developmental niche does not necessarily imply that its inclusion will improve an engineered microenvironment. Rather, our findings point to context-dependent interactions among extracellular matrix and soluble cues and support functional refinement of developmentally nominated factors rather than simple accumulation of niche-associated components^64^.

Our proteomic analysis of Matrigel further illustrates the distinction between a biologically effective matrix and a compositionally defined one. The three Matrigel lots examined in this study showed broadly similar proteomic profiles and produced comparable cellular responses under the conditions tested (**Figures S4, S5**). Notably, laminin-family proteins and type IV collagen were consistently abundant across all three lots, a finding that corroborates our human embryo scRNA-seq analysis, which prioritized laminin and collagen IV as candidate structural cues. This concordance provides additional support for the biological relevance of these matrix components in pluripotent cell culture. At the same time, more than 500 proteins were detected across the three lots, including a measurable fraction of non-matrisome and intracellular proteins. Thus, even when functional differences between lots are not apparent, the compositional complexity of Matrigel makes it difficult to assign specific cellular responses to individual matrix or signaling components^32,33^. By contrast, the defined ACLP formulation enables structural and soluble factors to be evaluated individually and in combination, facilitating mechanistic interpretation and providing greater control over bioink composition.

PTN provided a context-dependent supportive signal rather than a constitutive requirement for pluripotency (**Figures 3, 4**). Under standard E8 conditions, PTN supplementation had little additional effect, whereas under factor-deprived conditions it preserved cell viability, proliferation, and OCT4 expression. This behavior is consistent with earlier work showing that PTN supports human embryonic stem cell survival and expansion partly through PTPRZ1-dependent AKT activation^43^. In our bioprinted hiPSC system, PTPRZ1 knockdown similarly reduced AKT phosphorylation together with viability and pluripotency-associated responses, supporting a role for PTN–PTPRZ1 signaling under conditions in which exogenous pluripotency-supporting signals are limited. PTPRZ1 perturbation also reduced nuclear β-catenin and downstream Wnt-related transcription, suggesting crosstalk with pathways involved in pluripotent cell maintenance (**Figure S9**). Thus, PTN appears to function as a context-dependent supportive cue whose contribution becomes apparent when canonical pluripotency-supporting signals are reduced.

Beyond supporting pluripotent cell maintenance, the defined bioink also remained compatible with fabrication of distinct 3D geometries (**Figure 5**). This is relevant because bioprinting allows the biochemical composition of the cellular environment and the initial geometry of the construct to be controlled simultaneously. In this study, embryo-inspired geometries served as a proof of principle that niche composition and construct geometry can be controlled within the same defined platform. Although these structures were not intended to model embryogenesis, they establish a framework in which biochemical composition and prescribed starting geometry could be varied independently to examine their effects on subsequent cell organization, differentiation, and morphogenesis. Such an approach may be particularly useful for developmental biofabrication, where initial geometry can influence cell–cell interactions, diffusion fields, and mechanical boundary conditions.

Several limitations should nevertheless be acknowledged. First, although bioprinted hiPSCs retained multilineage developmental potential, including trilineage differentiation and teratoma formation **(Figure S7)**, long-term directed differentiation toward specific tissue lineages was not examined. Future studies should therefore determine whether the defined bioink supports sustained differentiation and maturation toward tissue-specific fates. Second, pluripotency and cell survival were assessed primarily over a 4-day culture period. Longer-term pluripotency and differentiation competence remain to be evaluated. Third, although our data implicate PTPRZ1 and AKT in the response to PTN, additional receptor-specific perturbation and rescue experiments will be required to define this signaling relationship more fully. Finally, PTN retention, release kinetics, and long-term bioactivity within the matrix were not directly measured. These parameters will be important to characterize if prolonged PTN-mediated support is required in future applications.

In conclusion, this study establishes an embryo-informed, transcriptomics-guided approach to designing a Matrigel-free bioink for undifferentiated hiPSC bioprinting. Single-cell analysis of the early human embryo identified extracellular and soluble niche cues that informed the development of a defined laminin/collagen IV-based bioink supporting hiPSC survival, proliferation, pluripotency, and multilineage developmental potential. PTN provided an additional supportive signal associated with PTPRZ1-dependent AKT signaling, particularly under conditions lacking exogenous pluripotency-supporting factors. Together, these findings show that selected functions of a complex, undefined matrix can be recapitulated using a minimal set of defined cues and establish developmental transcriptomic information as a biologically grounded framework for rational bioink design.

## Methods

### hiPSC culture and maintenance

Human induced pluripotent stem cells (hiPSCs; SCVI-111 line) derived from peripheral blood mononuclear cells were obtained from Stanford Cardiovascular Institute. hiPSCs were maintained on Matrigel-coated culture plates (Corning, Cat. No. 354230; 1:400 dilution in DMEM/F12; Welgene, Cat. No. LM002-04) in Essential 8 (E8) medium (Thermo Fisher Scientific, Cat. No. A1517001), with daily medium changes. Cells were passaged every 3–4 days using Ca²⁺/Mg²⁺-free DPBS (Welgene, Cat. No. LB001-02) containing 0.5 mM EDTA (Thermo Fisher Scientific, Cat. No. 15575020). For the first 24 h after passaging, E8 medium was supplemented with 10 μM ROCK inhibitor Y-27632 (BioGems, Cat. No. 1293823). Cells were maintained at 37 °C in a humidified incubator with 5% CO₂.

### Bioprinting

hiPSCs were dissociated into single cells by incubation with 0.5 mM EDTA in DPBS for 5 min, followed by gentle trituration. Cell concentration and viability were assessed using an automated cell counter (LUNA, Logos Biosystems). The cell suspension was then centrifuged at 300 × g for 4 min and resuspended at a concentration appropriate to achieve a final cell density of 100 × 10 cells/mL in the bioink.

A homogeneous bioink was prepared by mixing hiPSCs with 3% (w/v) alginate (Sigma-Aldrich, Cat. No. A2033) and either Matrigel or defined ECM components comprising laminin (Corning, Cat. No. 354259), collagen IV (Sigma-Aldrich, Cat. No. C5533), fibronectin (R&D Systems, Cat. No. 1918-FN-02M), and pleiotrophin (R&D Systems, Cat. No. 252-PL-050) as specified for each experimental condition. All bioink preparation steps were performed on ice.

The formulated bioink was loaded into a printing cartridge equipped with a 20-gauge (20G) nozzle and printed using a BIO X 3D bioprinter (CELLINK) at 5–20 kPa into 96-well plates (SPL Life Sciences, Cat. No. 30096).

Immediately following the printing process, the bioprinted droplets were crosslinked with 50 mM CaCl₂ (Sigma-Aldrich, Cat. No. C5670) in Ca²⁺/Mg²⁺-free DPBS for 5 min. The crosslinking solution was then removed, and the constructs were cultured in Essential 8 medium supplemented with 10 μM Y-27632.

### Bright-field imaging

Bright-field imaging of hiPSC-laden bioprinted constructs was performed daily from day 1 to day 4 using Cytation 5 and Cytation 10 Cell Imaging Multi-Mode Readers (BioTek Instruments) under consistent imaging settings. Images were acquired and analyzed using Gen5 software.

### Immunocytochemistry (ICC)

Immunofluorescence staining was performed to assess protein marker expression in hiPSC-laden 3D bioprinted droplets. Ca²⁺/Mg²⁺-containing DPBS (Welgene, Cat. No. LB001-01) was used throughout the staining procedure to maintain construct integrity. Constructs were fixed with 4% paraformaldehyde (PFA) (Thermo Fisher Scientific, Cat. No. 28908) prepared in Ca²⁺/Mg²⁺-containing DPBS overnight at 4 °C. Samples were subsequently washed five times for 30 min each with wash buffer consisting of 0.1% Tween-20 (Sigma-Aldrich, Cat. No. P1379) in Ca²⁺/Mg²⁺-containing DPBS. For permeabilization, samples were incubated with 0.1% Triton X-100 (Sigma-Aldrich, Cat. No. X100) in Ca²⁺/Mg²⁺-containing DPBS on a shaker for 1 h, and this step was repeated twice.

Samples were then incubated in blocking buffer containing 3% BSA, 2% goat serum (Abcam, Cat. No. ab7481), and 0.01% saponin (Sigma-Aldrich, Cat. No. 84510) in Ca²⁺/Mg²⁺-containing DPBS overnight at 4 °C.

Primary antibodies against OCT4 (Abcam, Cat. No. ab19857), TRA-1-60 (Abcam, Cat. No. ab16288), Ki67 (Abcam, Cat. No. ab16667), SOX17 (Thermo Fisher Scientific, Cat. No. MA5-24885), PAX6 (Thermo Fisher Scientific, Cat. No. MA1-109), Brachyury (Abcam, Cat. No. ab209665), PTPRZ1 (Abcam, Cat. No. ab126497), and β-catenin (Abcam, Cat. No. ab6302) were diluted 1:500 in blocking buffer and incubated with the samples for 24 h at 4 °C. Samples were washed five times with wash buffer for 30 min per wash.

Fluorophore-conjugated secondary antibodies including Alexa Fluor™ 488 goat anti-mouse IgG (H+L) (Thermo Fisher Scientific, Cat. No. A-11001), Alexa Fluor™ 568 goat anti-rabbit IgG (H+L) (Thermo Fisher Scientific, Cat. No. A-11011), Alexa Fluor™ 647 goat anti-mouse IgG (H+L) (Thermo Fisher Scientific, Cat. No. A-21235), and Alexa Fluor™ 647 goat anti-rabbit IgG (H+L), highly cross-adsorbed (Thermo Fisher Scientific, Cat. No. A-21245), were diluted 1:500 in blocking buffer and incubated for 2 h at room temperature in the dark. Hoechst 33342 (Thermo Fisher Scientific, Cat. No. H1399) was added during secondary antibody incubation for simultaneous nuclear counterstaining. Samples were then washed five times for 30 min each with wash buffer and mounted using mounting medium (Thermo Fisher Scientific, Cat. No. 00-4958-02). Fluorescence images were acquired using a confocal microscope.

### Quantitative Reverse Transcription PCR (qRT-PCR)

Total RNA was extracted using NucleoZOL reagent (Macherey-Nagel, Cat. No. MN740404.200) according to the manufacturer’s instructions. RNA was precipitated with 100% isopropanol (Sigma-Aldrich, Cat. No. I9526), washed with 75% ethanol, and resuspended in nuclease-free water (Thermo Fisher Scientific, Cat. No. AM9916). RNA concentration and purity were measured using a NanoDrop™ 2000 Spectrophotometer (Thermo Fisher Scientific), based on A260/A280 and A260/A230 ratios.

Complementary DNA (cDNA) was synthesized from 1 μg of total RNA using the High-Capacity cDNA Reverse Transcription Kit (Thermo Fisher Scientific, Cat. No. 4374967) in a final reaction volume of 20 μL according to the manufacturer’s instructions.

Quantitative real-time PCR was performed using cDNA, gene-specific primers, nuclease-free water, and KAPA SYBR® FAST qPCR Master Mix (Kapa Biosystems, Cat. No. KK4602) using a CFX Duet Real-Time PCR system (Bio-Rad). Reactions were performed in technical triplicates under the following cycling conditions, consisting of initial denaturation at 95 °C for 3 min, followed by 40 cycles of 95 °C for 5 s and 60 °C for 20 s. A melt curve analysis was subsequently performed from 65 °C to 95 °C in 0.5 °C increments. Relative gene expression was normalized to GAPDH and calculated using the 2^-ΔΔCT method. Primer sequences were obtained from the PrimerBank and synthesized by Cosmogenetech (Seoul, Republic of Korea).

### Conventional PCR

Conventional PCR was performed using cDNA templates synthesized as described above. PCR reactions were prepared using Phusion™ Plus PCR Master Mix (Thermo Fisher Scientific, Cat. No. F631L), gene-specific forward and reverse primers, nuclease-free water, and cDNA template. Forward and reverse primers were used at a final concentration of 0.5 μM. PCR amplification was performed using a thermal cycler under the following conditions: initial denaturation at 98 °C for 30 s, followed by 30 cycles of denaturation at 98 °C for 10 s, annealing at 60 °C for 10 s, and extension at 72 °C for 30 s, followed by a final extension at 72 °C for 5 min.

PCR products were separated by electrophoresis on a 2% agarose (Bio-Rad, Cat. No. 1613101) gel containing GelRed® Nucleic Acid Gel Stain (Biotium, Cat. No. 41003).

### Confocal microscopy

Immunofluorescently labeled samples were imaged using a STELLARIS 5 confocal microscope (Leica Microsystems). Image acquisition was performed utilizing LAS X software. Images were acquired using 10X, 20X, and 40X objective lenses, depending on the required field of view and resolution. Excitation and detection settings were optimized for the respective fluorophores. For quantitative comparisons, identical imaging settings, including laser power, pinhole size, and detector gain, were maintained across experimental groups. Z-stack images were acquired to evaluate the three-dimensional organization of the bioprinted constructs.

### Metabolic activity assay

Metabolic activity of hiPSCs within the 3D bioprinted constructs was assessed on days 1, 2, 3, and 4 after printing using the PrestoBlue™ Cell Viability Reagent (Thermo Fisher Scientific, Cat. No. A13261). PrestoBlue™ reagent was diluted at a 1:20 ratio in Essential 8 (E8) medium, and 200 μL of the working solution was added to each well containing the constructs. Following incubation for 3 h at 37 °C, absorbance was measured at 570 nm with 600 nm as the reference wavelength using a Cytation 5 Cell Imaging Multi-Mode Reader (BioTek Instruments). Data were acquired and analyzed using Gen5 software.

### Cell Viability Assay

Cell viability within the 3D bioprinted constructs was assessed using the LIVE/DEAD™ Viability/Cytotoxicity Kit (Thermo Fisher Scientific, Cat. No. L3224). A staining solution containing 2 μL of ethidium homodimer-1 (EthD-1) and 0.5 μL of calcein-AM per 1 mL of Essential 8 medium was prepared, and 100 μL was added to each well containing the bioprinted constructs. Samples were incubated for 10 min at 37 °C in the dark and imaged using a Cytation 5 Cell Imaging Multi-Mode Reader (BioTek Instruments). Images were acquired and analyzed using Gen5 software. Images used for viability quantification were acquired using a 4× objective. Cell viability (%) was calculated as the number of live cells divided by the total number of live and dead cells using ImageJ software.

### In Vitro Trilineage Differentiation

Following bioprinting, hiPSC-laden constructs were cultured in Essential 8 (E8) medium supplemented with 10 μM Y-27632 for 24 h. Differentiation toward the ectodermal, mesodermal, and endodermal lineages was then induced using the STEMdiff™ Trilineage Differentiation Kit (STEMCELL Technologies, Cat. No. 05230) according to the manufacturer’s protocol. Constructs were cultured in the corresponding lineage-specific differentiation medium with daily medium changes. Mesodermal and endodermal differentiation was performed for 5 days, whereas ectodermal differentiation was performed for 7 days. Following differentiation, lineage-specific marker expression was assessed by immunocytochemistry for PAX6 (ectoderm), Brachyury (mesoderm), and SOX17 (endoderm).

### Small Interfering RNA (siRNA) Transfection

hiPSCs were transfected with a PTPRZ1-targeting siRNA (Bioneer) or Silencer™ Select Negative Control No. 1 siRNA (Thermo Fisher Scientific, Cat. No. 4390843). Cells were seeded in 12-well plates and cultured to approximately 60–70% confluence before transfection. siRNA transfection was performed using Lipofectamine™ RNAiMAX Transfection reagent (Thermo Fisher Scientific, Cat. No. 13778150). siRNA and RNAiMAX complexes were prepared in Opti-MEM™ | Reduced Serum Medium (Thermo Fisher Scientific, Cat. No. 31985062) according to the manufacturer’s instructions and added to cells at a final siRNA concentration of 50 nM. After 48 h, cells were harvested, and PTPRZ1 knockdown was assessed by qRT-PCR and Western blotting.

### Western blot

Total protein was extracted using RIPA Lysis and Extraction Buffer (Thermo Fisher Scientific, Cat. No. 89900) supplemented with Halt™ Protease and Phosphatase Inhibitor Cocktail (100X) (Thermo Fisher Scientific, Cat. No. 78440). Cell lysates were incubated on ice for 30 min and subsequently centrifuged at 13,000 × g for 10 min at 4 °C. The supernatant was collected, and protein concentrations were determined using the Pierce™ BCA Protein Assay Kit (Thermo Fisher Scientific, Cat. No. 23225).

Equal amounts of protein (5–10 μg/lane) were separated by 3–8% SDS-PAGE and transferred onto PVDF membranes. Membranes were blocked with 5% skim milk in TBST for 1 h at room temperature and incubated overnight at 4 °C with the following primary antibodies: PTPRZ1 (Abcam, Cat. No. ab126497), pan-Akt (Cell Signaling Technology, Cat. No. 4691), Phospho-Akt (Ser473) (Cell Signaling Technology, Cat. No. 4060), and GAPDH (Thermo Fisher Scientific, Cat. No. MA5-15738) at the indicated dilutions.

Following three washes with TBST, the membranes were incubated for 1 h at room temperature with HRP-conjugated secondary antibodies: Anti-rabbit IgG, HRP-linked antibody (Cell Signaling Technology, Cat. No. 7074P2) and anti-mouse IgG, HRP-linked antibody (Cell Signaling Technology, Cat. No. 7076P2). Protein bands were detected using an enhanced chemiluminescence reagent (Thermo Fisher Scientific, Cat. No. 34577) and imaged using a ChemiDoc Imaging System (Bio-Rad). Band intensities were quantified using ImageJ software.

### CAD design and FRESH-assisted bioprinting of embryo-inspired constructs

Three embryo-inspired constructs representing the inner cell mass (ICM), bilaminar disc, and primitive streak were designed using Autodesk Fusion (Autodesk, Inc.) and exported in STL format for printing. To support the fabrication of these complex three-dimensional geometries, a FRESH supporting bath was prepared.

The FRESH supporting bath consisted of a 1:1 mixture of absolute ethanol (Merck, Cat. No. 1.00983) and autoclaved distilled water containing 2% (w/v) gelatin type B (Sigma-Aldrich, Cat. No. G9391), 0.25% (w/v) Pluronic F-127 (Sigma-Aldrich, Cat. No. P2243), and 0.1% (w/v) gum Arabic (Sigma-Aldrich, Cat. No. G9752). The mixture was stirred continuously at 250 rpm overnight. The resulting suspension was transferred to 50-mL conical tubes and centrifuged at 300 × g for 5 min at room temperature. The supernatant was removed, and the FRESH material was washed with 10 mL of 10 mM CaCl₂ in DPBS, followed by centrifugation at 1,000 × g for 2 min at room temperature. The washing procedure was repeated three times. After the final wash, the supernatant was completely removed, and the FRESH supporting bath was used immediately for bioprinting. For acellular printing, the ACLP bioink without cells was printed into the FRESH supporting bath according to the corresponding STL designs. For cellular printing, hiPSCs were incorporated into the ACLP bioink at a final density of 100 × 10 cells/mL and printed using the same designs. Printing was performed using a BIO X 3D bioprinter (CELLINK) equipped with a 25-gauge nozzle at 5–20 kPa. Following printing, the constructs were incubated at 37 °C for 1 h to liquefy and remove the FRESH supporting bath. The constructs were then crosslinked with 50 mM CaCl₂ in Ca²⁺/Mg²⁺-free DPBS for 5 min, washed with Ca²⁺/Mg²⁺-containing DPBS, and subsequently transferred to culture medium. Cellular constructs were cultured in Essential 8 medium supplemented with 10 μM Y-27632 for the first 24 h after printing.

### Cryosectioning and immunofluorescence staining of printed constructs

Day-4 printed ICM constructs were fixed with 4% formaldehyde in Ca²⁺/Mg²⁺-containing DPBS overnight at 4 °C. Fixed constructs were embedded in Surgipath FSC 22 Clear Frozen Section Compound (Leica Biosystems, Cat. No. 3801480) and stored at −80 °C until cryosectioning. Constructs were sectioned at 50 or 100 μm thickness using a Microm HM 525 cryostat (Thermo Scientific) and mounted onto glass slides.

For immunofluorescence staining, a permeabilization/wash buffer consisting of 0.1% Triton X-100 in Ca²⁺/Mg²⁺-containing DPBS was used throughout the staining procedure. Cryosections were equilibrated at room temperature for approximately 30 min and permeabilized with the permeabilization/wash buffer for 20 min at room temperature. Sections were then blocked with 5% goat serum prepared in permeabilization/wash buffer for 1 h at room temperature.

Primary antibodies against OCT4 (Abcam, Cat. No. ab19857) and TRA-1-60 (Abcam, Cat. No. ab16288) were diluted 1:1000 in 2.5% goat serum prepared in permeabilization/wash buffer and incubated with the sections overnight at 4 °C.

The following day, sections were washed three times for 5 min each with permeabilization/wash buffer. Fluorophore-conjugated secondary antibodies were diluted 1:1000 in 2.5% goat serum, and DAPI was added simultaneously at a 1:1000 dilution. Sections were incubated for 1 h at room temperature in the dark and subsequently washed three times for 10 min each with permeabilization/wash buffer.

Sections were briefly rinsed with distilled water, air-dried for approximately 1 h, mounted using mounting medium, and allowed to set for approximately 2 h before storage at 4 °C.

### Printing fidelity and structural stability analysis

Printing fidelity was evaluated by comparing the dimensions of printed constructs with the corresponding CAD-defined dimensions. Top- and side-view images of the inner cell mass (ICM), bilaminar disc, and primitive streak constructs were acquired, and dimensions were measured using ImageJ software. Image scale was calibrated using the scale bar included in each image. Dimensional fidelity was calculated as:

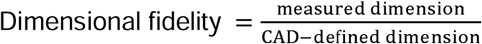

To assess structural stability during culture, ICM constructs were imaged on days 1, 2, and 4 after printing. Dimensions were measured using ImageJ and normalized to the corresponding day-1 value:

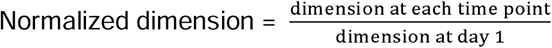

### Single-cell RNA-seq data

#### Pre-processing

Publicly available human embryonic single-cell RNA sequencing (scRNA-seq) datasets from Xiang et al. (GEO: GSE136447), Molè et al. (ArrayExpress: E-MTAB-8060), and Zhou et al. (GEO: GSE109555) were used for the analysis^34–36^. The datasets collectively covered human embryonic developmental stages from 6 to 14 days post-fertilization (DPF). For the Xiang dataset, gene identifiers were converted to gene symbols, and duplicated gene symbols were consolidated by summing their expression values.

Quality control was performed independently for each dataset. Cells from the Xiang dataset with 7,800–15,800 detected genes and <30% mitochondrial transcripts were retained. Cells from the Zhou dataset with 2,300–11,900 detected genes were retained, whereas cells from the Molè dataset with >5,000 detected genes and <30% mitochondrial transcripts were retained for downstream analysis.

#### Data analysis

Each dataset was independently normalized using the global-scaling normalization method implemented in NormalizeData in Seurat, followed by identification of 2,000 highly variable features using the variance-stabilizing transformation (VST) method. The datasets were subsequently integrated using the FastMNN method implemented in SeuratWrappers to correct dataset-specific batch effects^65^.

Using the calculated MNN reduction, dimensionality reduction was performed using uniform manifold approximation and projection (UMAP). A nearest-neighbor graph was constructed using the first 30 MNN dimensions, followed by graph-based clustering at a resolution of 0.08.

Cell populations were annotated based on the expression of lineage-associated marker genes, including ARGFX and VENTX for epiblast, DPF3 and CDX2 for inner cell mass, HES2 and FOSL1 for syncytiotrophoblasts, PDGFRA and SOX17 for primitive endoderm, ASCL2 for extravillous trophoblasts, and NR2F2 for cytotrophoblasts. The annotated epiblast population was subsequently selected for downstream ligand–receptor interaction analysis.

#### Ligand-Receptor interaction and candidate filtering analysis

The annotated integrated scRNA-seq dataset was stratified into pre-implantation (6–10 DPF) and post-implantation (9–14 DPF) developmental groups based on the original developmental-stage metadata. Ligand–receptor interactions were analyzed separately for each group using the LIANA (LIgand-receptor ANalysis frAmework) R package, which integrates multiple cell–cell communication inference methods^66^. Interactions were inferred using liana_wrap() and subsequently aggregated using liana_aggregate() to obtain consensus-ranked ligand–receptor interactions.

Primitive endoderm (PrE), epiblast (EPI), inner cell mass (ICM), cytotrophoblasts (CTBs), syncytiotrophoblasts (STBs), and extravillous trophoblasts (EVTs) were included as ligand-producing source populations, whereas EPI was designated as the target population. Epiblast-targeted ligand–receptor interactions were ranked according to the LIANA consensus results, and the top 25 ligand candidates from each developmental group were selected for subsequent analysis. Ligands identified in both developmental groups were classified as conserved, whereas those preferentially identified in either group were classified as stage-specific.

Receptor genes corresponding to the selected ligand–receptor interactions were submitted to the Database for Annotation, Visualization and Integrated Discovery (DAVID) for KEGG pathway enrichment analysis^67^. All KEGG pathways identified from the receptor gene set were initially considered, and pathways associated with pluripotency maintenance and related signaling processes were selected for subsequent candidate prioritization. Ligands associated with these pathways were further evaluated based on previously reported roles in pluripotent stem-cell maintenance, self-renewal, proliferation, survival, and cell–matrix interactions.

### Statistical analysis

Statistical analysis was performed using GraphPad Prism 10. Data are expressed as mean ± SEM from at least three independent biological replicates, unless otherwise indicated. Normality was assessed prior to applying t-tests or one/two-way ANOVA, with appropriate post-hoc corrections. Statistical significance was set at **p < 0.05** (∗p < 0.05, ∗∗p < 0.01, ∗∗∗p < 0.001, ∗∗∗∗p < 0.0001).

### Proteomics

#### Preparation of Matrigel samples for proteomic analysis

Three independent lots of Matrigel (Corning, Cat. No. 354230; Lot Nos. 27624003, 16925001, and 26625004) were analyzed to assess lot-to-lot variability in protein composition. Sample preparation was performed using the Filter-Aided Sample Prep Protein Digestion Kit (Abcam, Cat. No. ab270519) with modifications to the manufacturer’s protocol. Matrigel protein samples were diluted in 50 mM ammonium bicarbonate buffer, followed by disulfide bond reduction with DL-dithiothreitol (Sigma-Aldrich, Cat. No. D0632) at a final concentration of 10 mM. Reduction was performed for 30 min at 60 °C in a heating block. Cysteine residues in the protein samples were alkylated with iodoacetamide (IAA) at a final concentration of 55 mM for 30 min at room temperature under light-protected conditions. Proteolytic digestion was then carried out using proteomics-grade trypsin (Sigma-Aldrich, Cat. No. T6567) at an enzyme-to-protein ratio of 1:50 for 16 h at 37 °C.

Following digestion, peptides were purified using Pierce™ Peptide Desalting Spin Columns (Thermo Fisher Scientific, Cat. No. 89851), according to the manufacturer’s protocol, and the peptides retained on the column were recovered with an elution solution containing 0.1% (v/v) liquid chromatography-mass spectrometry (LC-MS)-grade formic acid (Tokyo Chemical Industry, Cat. No. F0654) and 50% (v/v) high-performance liquid chromatography (HPLC)-grade acetonitrile (Thermo Fisher Scientific, Cat. No. A998SK). The purified peptides were dried and concentrated using a centrifugal evaporator (CVE-3000; Eyela, Tokyo, Japan). Dried peptides were reconstituted in 0.1% (v/v) formic acid containing 2% (v/v) acetonitrile. Peptide concentrations were quantified using the Pierce™ Quantitative Colorimetric Peptide Assay (Thermo Fisher Scientific, Cat. No. 23275) according to the manufacturer’s instructions, with absorbance measured at 480 nm using an Infinite M200 Pro microplate reader (Tecan, Männedorf, Switzerland). All samples were adjusted to a final concentration of 0.5 mg mL^-1^ for downstream analysis.

#### Liquid chromatography tandem-mass spectrometry (LC-MS/MS)

For LC-MS/MS analysis, 2 µg of each peptide sample was prepared in a solvent containing 0.1% (v/v) formic acid and 2% (v/v) acetonitrile, and analyzed using an Orbitrap Exploris 240 mass spectrometer (Thermo Fisher Scientific) coupled to an UltiMate 3000 RSLCnano system (Thermo Fisher Scientific). Mobile phase A consisted of 0.1% (v/v) formic acid and 2% (v/v) acetonitrile, and mobile phase B consisted of 0.1% (v/v) formic acid in acetonitrile. Peptide samples were injected into an Acclaim™ PepMap™ 100 C18 trap column (Thermo Fisher Scientific, Cat. No. 164564) and washed with mobile phase A at a flow rate of 4 µL min^-1^ for 6 min. Peptides were subsequently separated on an EASY-Spray HPLC analytical column (Thermo Fisher Scientific, Cat. No. ES903). The peptide separation was performed at a flow rate of 250 nL min^-1^, and the gradient of mobile phase B was programmed as follows: 2%–8% over 10 min, 8%–30% over 55 min, 30%–90% over 1 min, maintenance at 90% for 4 min, 2% for 20 min. Mass spectrometric (MS) data were recorded using Xcalibur data acquisition software (version 4.5; Thermo Fisher Scientific). Full-scan MS spectra were collected in the Orbitrap mass analyzer with positive polarity across a mass-to-charge ratio (m/z) range of 350–1,800 at a resolution of 60,000 with a normalized automatic gain control (AGC) of 300%. For MS/MS acquisition, precursor ions were fragmented by higher energy collision dissociation (HCD) using a collision energy of 30%. MS/MS spectra were acquired at a resolution of 15,000 with a normalized AGC of 100% and maximum injection of 100 ms.

#### Proteomic data processing and analysis

Raw LC-MS/MS data were processed using MaxQuant^68,69^ (version 2.7.5; Max Planck Institute of Biochemistry, Munich, Germany). Protein identification was performed against the Mus musculus reference proteome from the UniProt database^70^ (released in January 2026). Data processing was conducted using the default MaxQuant parameter settings with trypsin specified as the digestion enzyme. The false discovery rate (FDR) was set to 1% at both the peptide and protein levels, and the “Match between runs” feature was enabled to facilitate consistent protein identification across LC-MS/MS runs. Protein abundances were determined using the intensity-based absolute quantification (iBAQ) algorithm, and the resulting iBAQ values were used for the subsequent data analyses.

Comparative analysis of protein profiles and abundances across three Matrigel lots was performed to characterize lot-to-lot variability in protein composition. For matrisome annotation and classification, the identified proteins were categorized as matrisome or non-matrisome proteins based on the Mus musculus matrisome database (rev. 2014) provided by the MIT Matrisome Project^71^. Matrisome proteins identified in each Matrigel lot were further classified into core matrisome proteins and matrisome-associated proteins. Specifically, core matrisome proteins comprised collagens, glycoproteins, and proteoglycans, whereas matrisome-associated proteins encompassed ECM-affiliated proteins, ECM regulators, and secreted factors.

### Teratoma assay

#### Teratoma formation

A testicular teratoma formation assay was performed to evaluate the in vivo differentiation potential of hiPSCs following bioprinting. Male BALB/c nude mice were obtained from ORIENTBIO at 4 weeks of age and housed individually under controlled temperature and humidity. Following a 2-week acclimatization period, cell transplantation was performed at 6 weeks of age. All animal procedures were approved by the Institutional Animal Care and Use Committee of Chung-Ang University (Approval No. 202501020157). Before transplantation, cell number and viability were assessed using a trypan blue exclusion assay. The mean cell viability was 93.0%. For each mouse, 3 × 10 viable cells were suspended in 30 μL of DPBS without Matrigel and injected into the right testis under isoflurane inhalation anesthesia. Teratomas were harvested 16 weeks after transplantation.

#### Histological analysis

Teratoma tissues were trimmed, fixed in 4% PFA for 48 h at 4 °C, dehydrated in graded ethanol, cleared in xylene and infiltrated with paraffin before embedding. Paraffin sections (5 μm) were deparaffinized and rehydrated before hematoxylin and eosin (H&E) staining using the H&E Staining Kit (Abcam, Cat. No. ab245880). Hematoxylin and eosin staining times were 5 min and 1–1.5 min, respectively. Additional sections were stained with Masson’s trichrome (MT) using the Trichrome Stain Kit (Abcam, Cat. No. ab150686). H&E was used to assess tissue morphology, and MT provided complementary visualization of extracellular matrix and muscle. Stained sections were dehydrated, cleared in xylene, mounted and examined using an microscope (Nikon). Sections were evaluated morphologically for derivatives of the three embryonic germ layers, including neural rosette-like structures (ectoderm), cartilage and muscle (mesoderm), and gut-like and secretory epithelium (endoderm).

### Ethics statement

This study was reviewed by the Institutional Review Board of Sungkyunkwan University and was determined to qualify for exemption from IRB review (File No. 2026-06-050). No new human participants were recruited, and no human biological specimens were collected in this study. The experiments were performed using an established human induced pluripotent stem cell (hiPSC) line.

### Declaration of AI-assisted language editing

Generative AI tools were used solely to improve the clarity, grammar, and readability of the manuscript. All scientific content, data interpretation, and conclusions were developed and verified by the authors. The authors reviewed and approved all AI-assisted edits and take full responsibility for the final manuscript.

## Data availability

The publicly available human embryonic single-cell RNA-sequencing datasets analyzed in this study are available from the Gene Expression Omnibus (GEO) under accession codes GSE136447 and GSE109555, and from ArrayExpress under accession code E-MTAB-8060. Source data underlying the graphs and charts presented in the main and supplementary figures will be provided with the paper.

## Acknowledgements

We thank the Stanford Cardiovascular Institute for providing the SCVI-111 human induced pluripotent stem cell (hiPSC) line used in this study. Figures were created using BioRender. Created in BioRender. Lee, S. (2026) https://BioRender.com/ft4rxaa.

## Author contributions

G.J., B.K. and S.L. conceived and designed the study. G.J. and J.K. performed the single-cell RNA-seq and bioinformatic analyses. G.J. and B.K. performed hiPSC culture, bioink formulation, 3D bioprinting and cellular assays. G.J. performed the mechanistic experiments, analyzed the data and prepared the figures. M.J.L. performed and analyzed the proteomic experiments under the supervision of S.W.C. J.C. and H.S.Y. performed and analyzed the teratoma assays under the supervision of S.H.M. G.J. and S.L. wrote the original draft and revised the manuscript. S.L. provided resources, supervised the study, administered the project and acquired funding. All authors reviewed and approved the final manuscript.

