## Supplementary Information for "Human embryo-informed niche design enables defined bioprinting of undifferentiated hiPSCs"

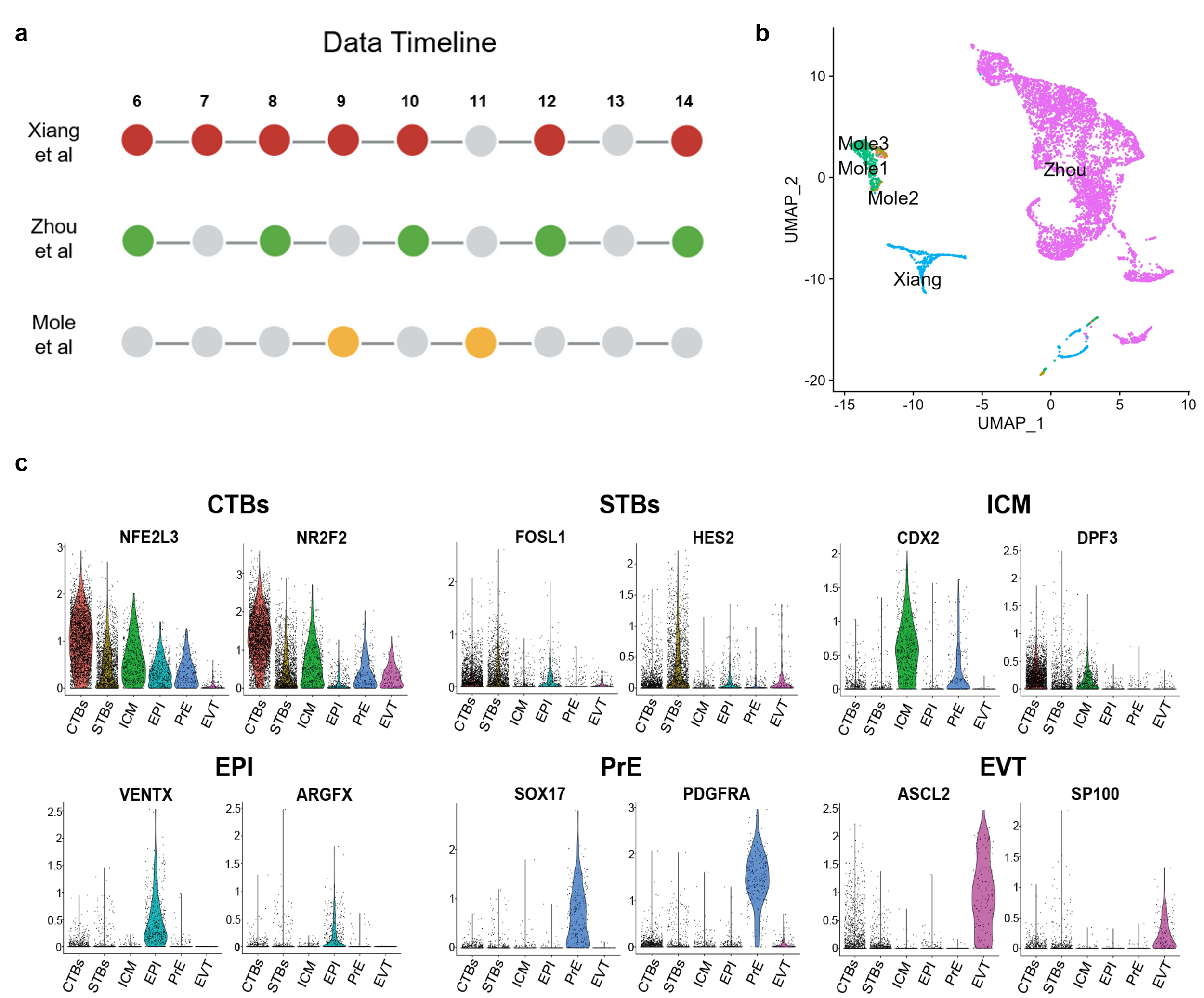

**Supplementary Fig. 1 | Overview and marker-based annotation of integrated public human embryo scRNA-seq datasets.**

**a,** Developmental-stage distribution of the three public human embryo scRNA-seq datasets from Xiang *et al.*, Zhou *et al.* and Mole *et al.*, covering post-fertilization days 6–14. Colored circles indicate developmental stages represented in each dataset, whereas gray circles indicate stages not included. **b,** UMAP visualization of the integrated datasets before batch correction, showing the distribution of cells according to their dataset of origin. **c,** Violin plots showing the expression of representative lineage-marker genes used for cell-type annotation, including cytotrophoblasts (CTBs; *NFE2L3*, *NR2F2*), syncytiotrophoblasts (STBs; *FOSL1*, *HES2*), inner cell mass (ICM; *CDX2*, *DPF3*), epiblast (EPI; *VENTX*, *ARGFX*), primitive endoderm (PrE; *SOX17*, *PDGFRA*) and extravillous trophoblasts (EVTs; *ASCL2*, *SP100*).

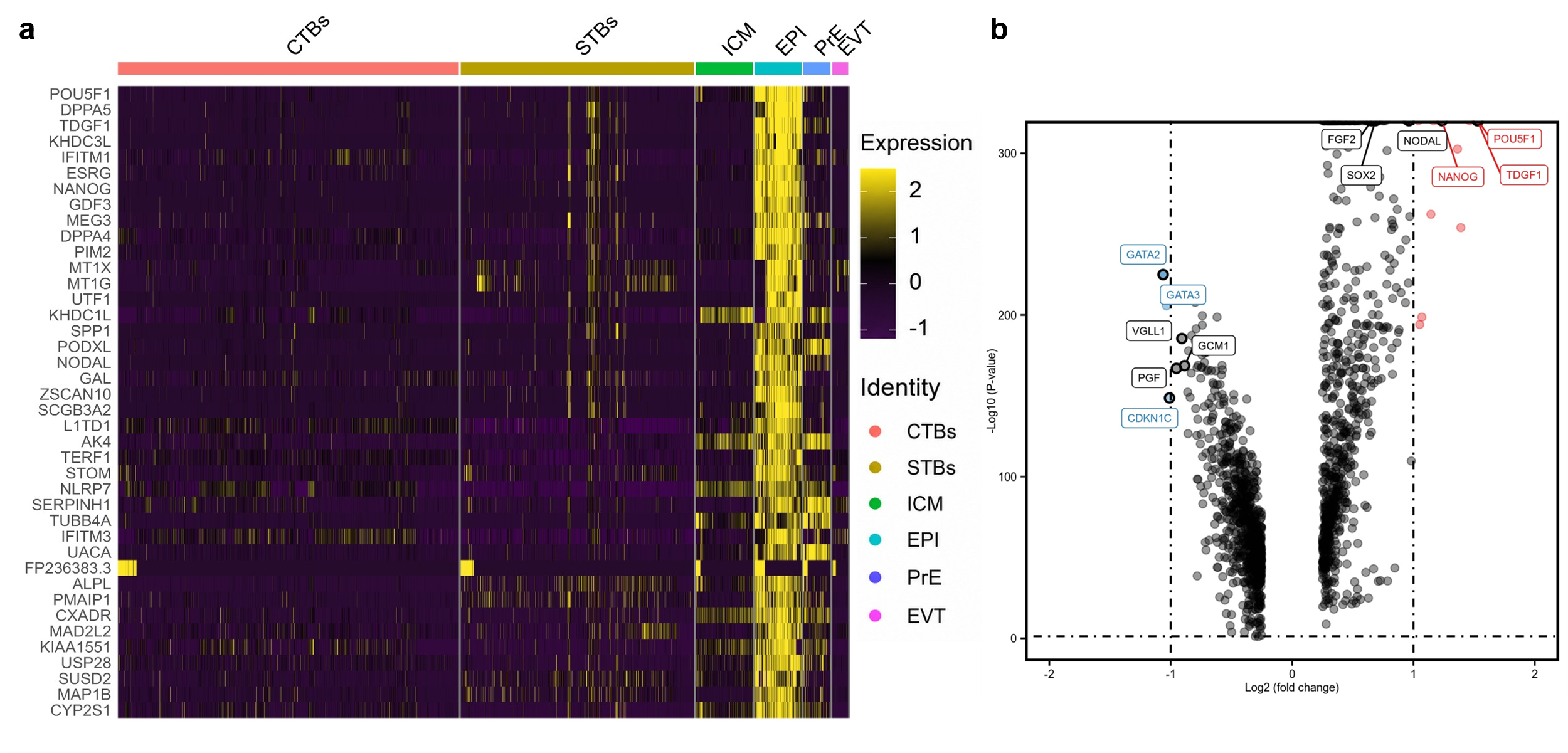

**Supplementary Fig. 2 | Transcriptomic characterization of the EPI cluster in the integrated in vivo human embryo scRNA-seq dataset.**

**a,** Heatmap showing the expression of representative lineage- and pluripotency-associated genes across annotated cell populations, including cytotrophoblasts (CTBs), syncytiotrophoblasts (STBs), inner cell mass (ICM), epiblast (EPI), primitive endoderm (PrE) and extravillous trophoblasts (EVTs). Pluripotency-associated genes are preferentially enriched in the EPI population. **b,** Volcano plot showing differentially expressed genes in EPI cells relative to the other annotated cell populations. Positive log2 fold-change values indicate genes enriched in EPI cells, including core pluripotency-associated markers such as *POU5F1*, *NANOG*, *TDGF1* and *NODAL*, whereas negative values indicate genes enriched in non-EPI populations. Dotted lines indicate the thresholds used for differential-expression analysis.

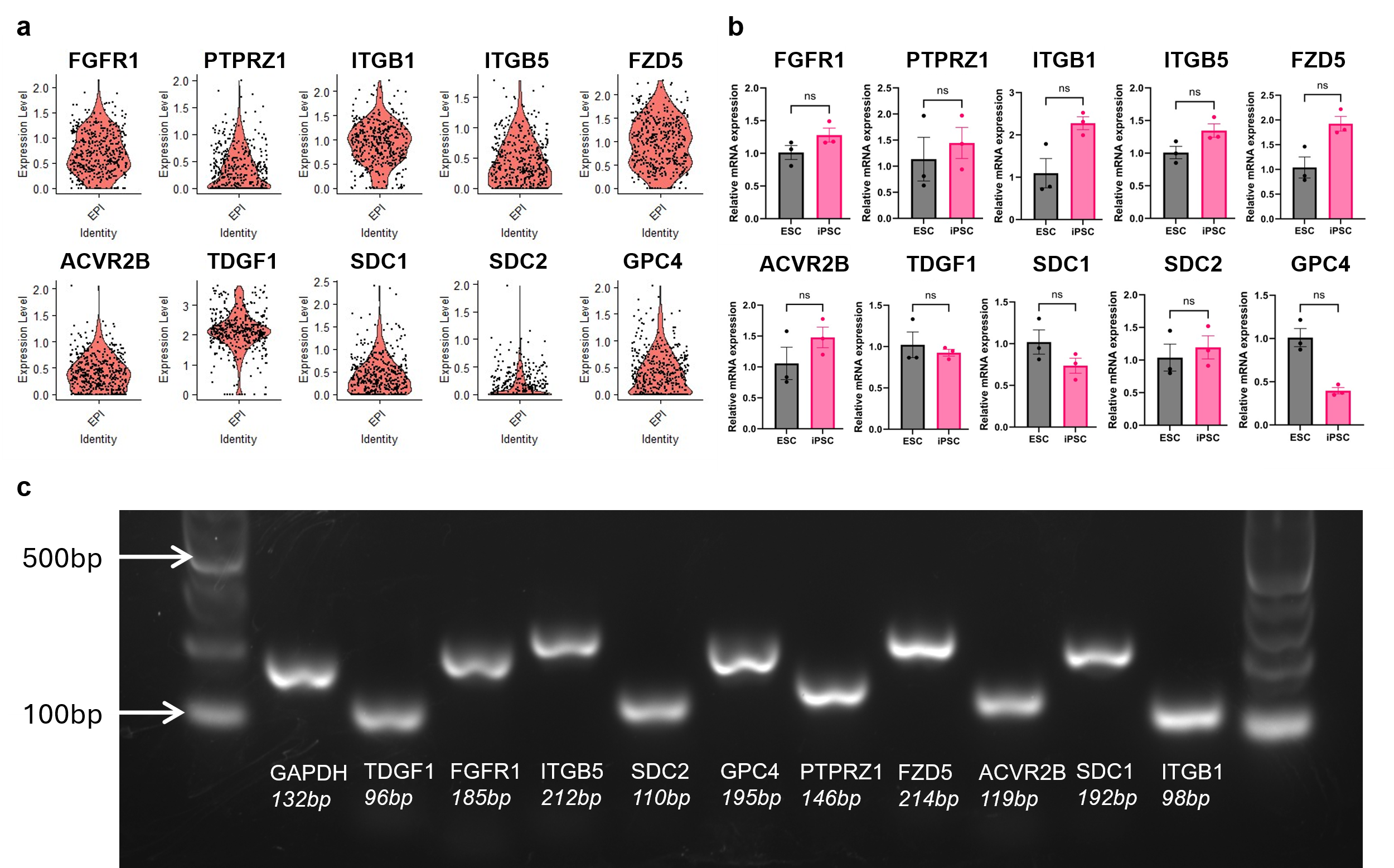

**Supplementary Fig. 3 | Validation of candidate receptor expression in in vivo epiblast cells and in vitro pluripotent stem cells.**

**a,** Violin plots showing the expression of candidate receptor genes in the EPI cluster of the integrated in vivo human embryo scRNA-seq dataset, including FGFR1, PTPRZ1, ITGB1, ITGB5, FZD5, ACVR2B, TDGF1, SDC1, SDC2 and GPC4. **b,** qRT–PCR analysis comparing the expression of the candidate receptor genes between ESCs and iPSCs. ns, not significant. **c,** Conventional RT–PCR analysis of candidate receptor expression in iPSCs, with PCR products resolved by agarose gel electrophoresis. GAPDH was used as an internal control, and the expected amplicon sizes are indicated below each gene.

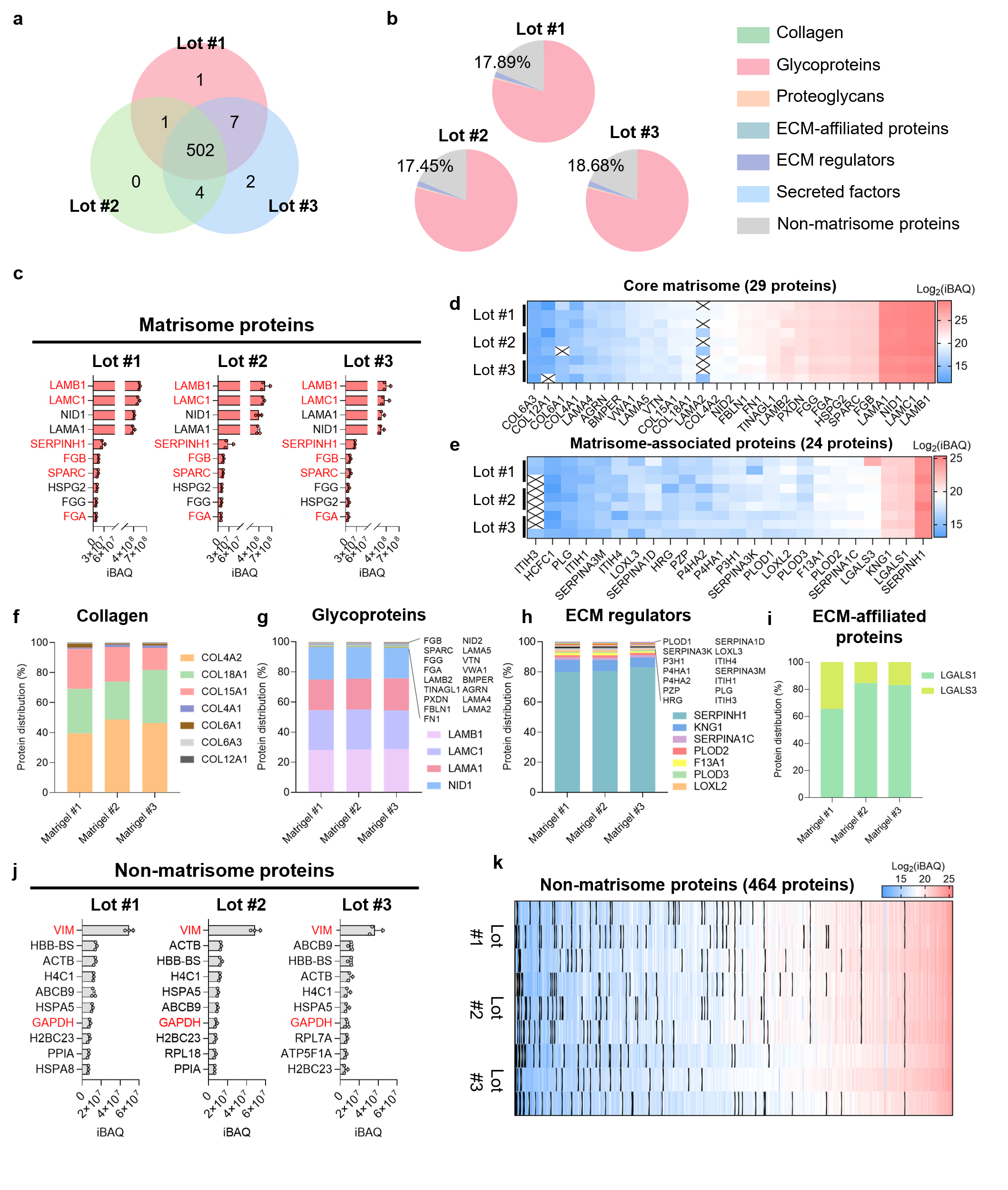

**Supplementary Fig. 4 | Proteomic characterization of Matrigel across independent lots.**
**a,** Venn diagram showing the overlap of proteins identified by mass spectrometry across three independent Matrigel lots. A total of 502 proteins were commonly detected in all three lots. **b,** Distribution of identified proteins according to matrisome classification in each Matrigel lot, including collagens, glycoproteins, proteoglycans, ECM-affiliated proteins, ECM regulators, secreted factors and non-matrisome proteins. **c,** Abundance of representative matrisome proteins in each Matrigel lot, quantified by intensity-based absolute quantification (iBAQ). **d,** Heatmap showing the abundance of 29 core matrisome proteins across the three Matrigel lots. **e,** Heatmap showing the abundance of 24 matrisome-associated proteins across the three lots. Protein abundance in d, e is presented as log2-transformed iBAQ values. **f–i,** Relative protein distribution within major matrisome categories across the three Matrigel lots: f, collagens; g, glycoproteins; h, ECM regulators; and i, ECM-affiliated proteins. **j,** Abundance of representative non-matrisome proteins identified in each Matrigel lot, quantified by iBAQ. **k,** Heatmap showing the abundance profiles of 464 non-matrisome proteins across the three Matrigel lots, presented as log2-transformed iBAQ values.

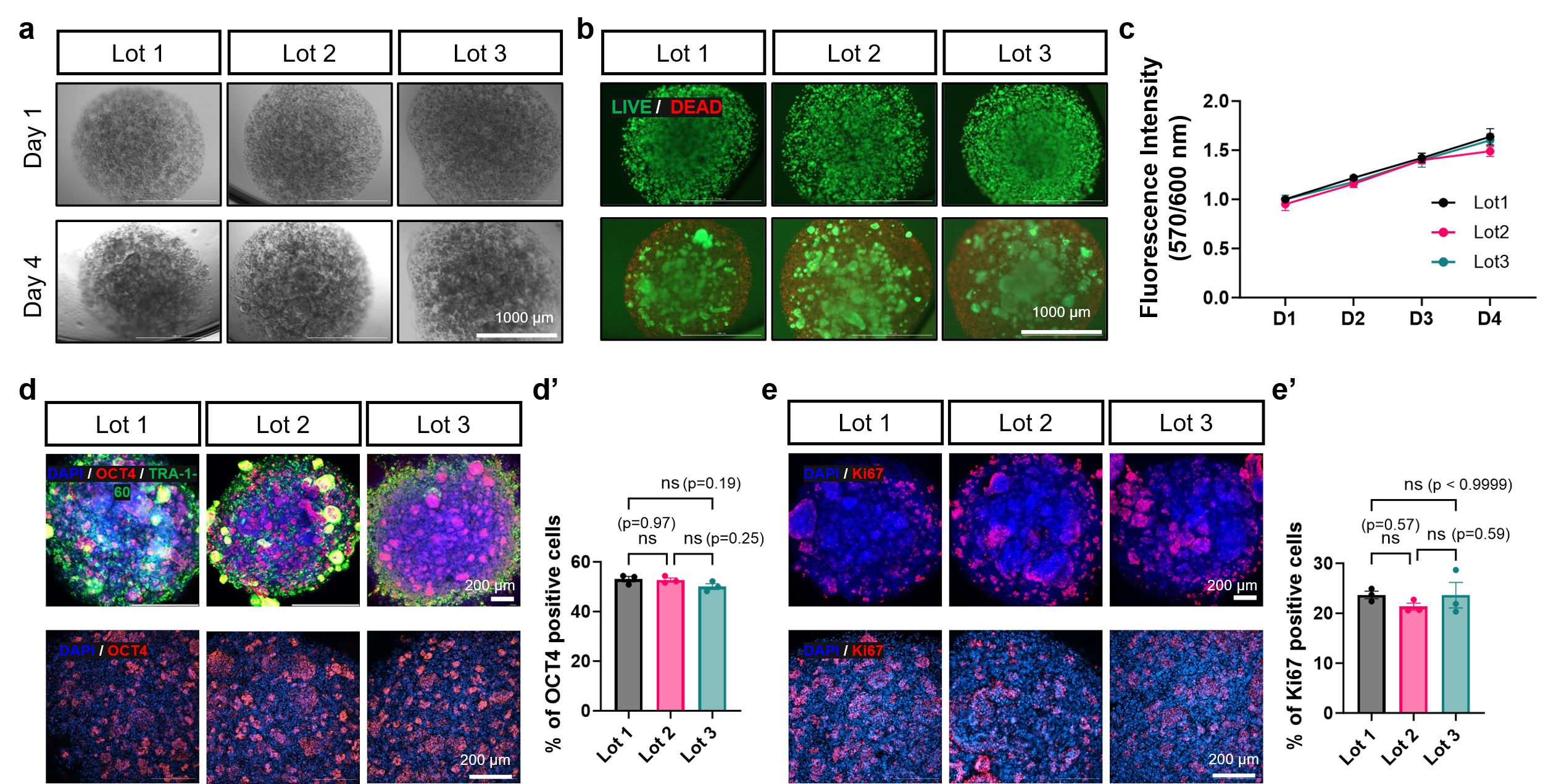

**Supplementary Fig. 5 | Biological performance of hiPSC constructs across different Matrigel lots.**
**a,** Representative bright-field images of hiPSC constructs generated using three independent Matrigel lots at the indicated culture time points. Scale bar, 1000 μm. **b,** Representative Live/Dead fluorescence images of constructs generated using the three Matrigel lots, showing viable (green) and dead (red) cells. Scale bar, 1000 μm. **c,** Quantification of cell viability over the culture period across the three Matrigel lots. **d,** Representative immunofluorescence images showing expression of the pluripotency markers OCT4 and TRA-1-60 in constructs generated using the three Matrigel lots. Nuclei were counterstained with DAPI. Scale bar, 200 μm. **d′,** Quantification of pluripotency-associated marker expression across the three Matrigel lots. **e,** Representative immunofluorescence images showing expression of the proliferation marker Ki67 in constructs generated using the three Matrigel lots. Nuclei were counterstained with DAPI. Scale bar, 200 μm. **e′,** Quantification of Ki67-positive cells across the three Matrigel lots. No significant lot-dependent differences were observed in the measured viability, pluripotency or proliferative responses.

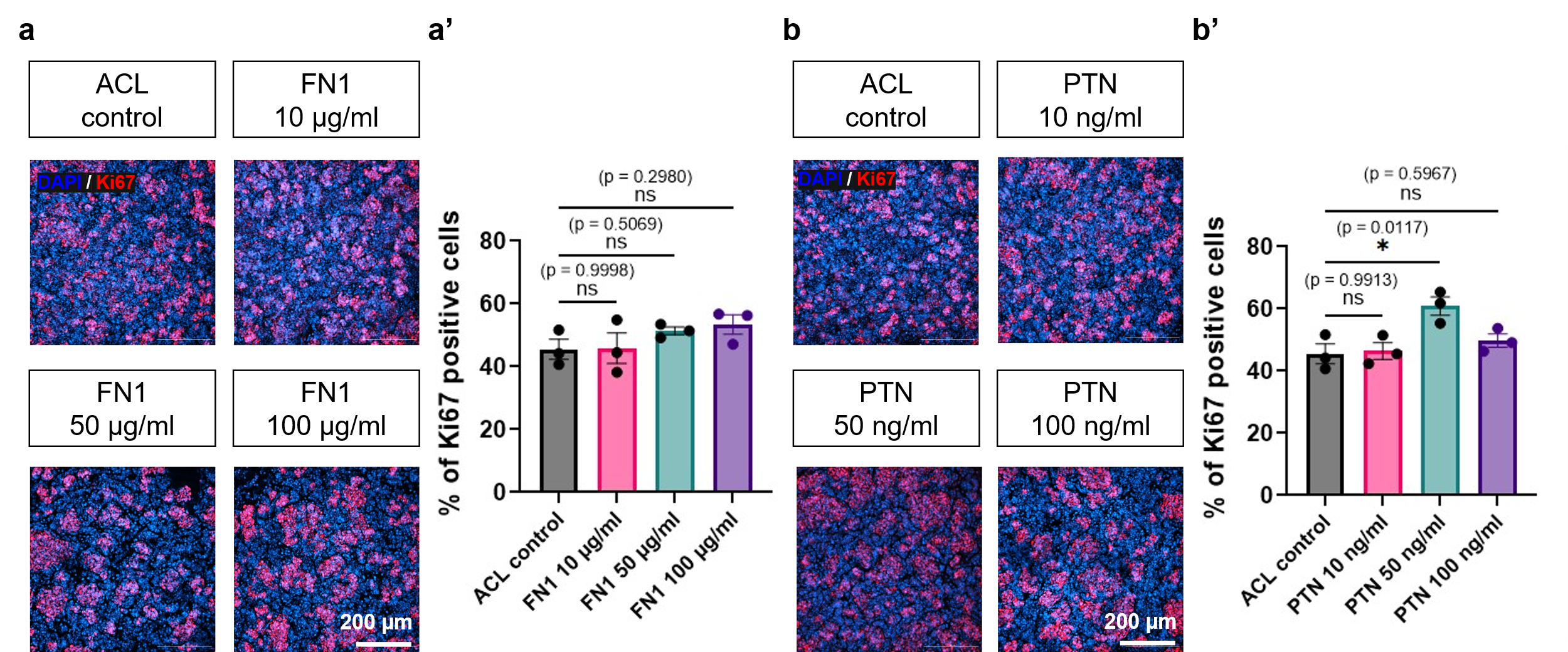

**Supplementary Fig. 6 | Optimization of FN1 and PTN supplementation in biomimetic bioinks.**
**a,** Representative immunofluorescence images of Ki67 expression in hiPSC constructs supplemented with FN1 at 10, 50 or 100 μg ml⁻¹. Nuclei were counterstained with DAPI. Scale bars, 200 μm. **a′,** Quantification of Ki67-positive cells in the ACL control and FN1-supplemented groups. **b,** Representative immunofluorescence images of Ki67 expression in hiPSC constructs supplemented with PTN at 10, 50 or 100 ng ml⁻¹. Nuclei were counterstained with DAPI. Scale bars, 200 μm. **b′,** Quantification of Ki67-positive cells in the ACL control and PTN-supplemented groups. FN1 at 50 μg ml⁻¹ and PTN at 50 ng ml⁻¹ showed the highest mean Ki67-positive fractions and were selected for subsequent experiments. Data are presented as mean ± SEM. Exact *P* values are indicated in the graphs; ns, not significant.

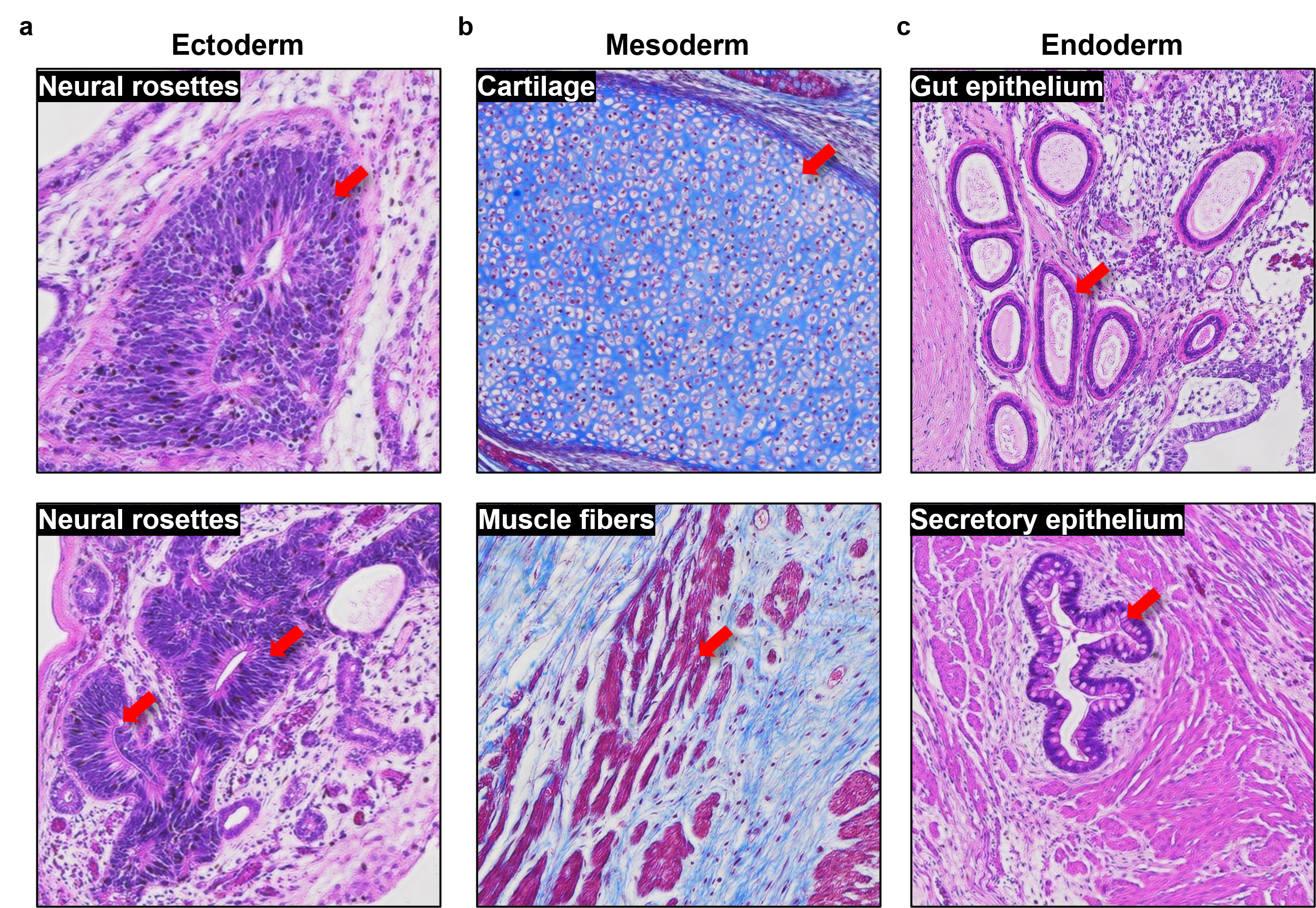

**Supplementary Fig. 7 | Histological validation of trilineage differentiation potential in iPSC-derived teratomas.**
**a,** Representative neural rosette structures indicating ectodermal differentiation. **b,** Representative cartilage and muscle fibers indicating mesodermal differentiation. **c,** Representative gut epithelium and secretory epithelium indicating endodermal differentiation. Red arrows indicate representative lineage-associated structures.

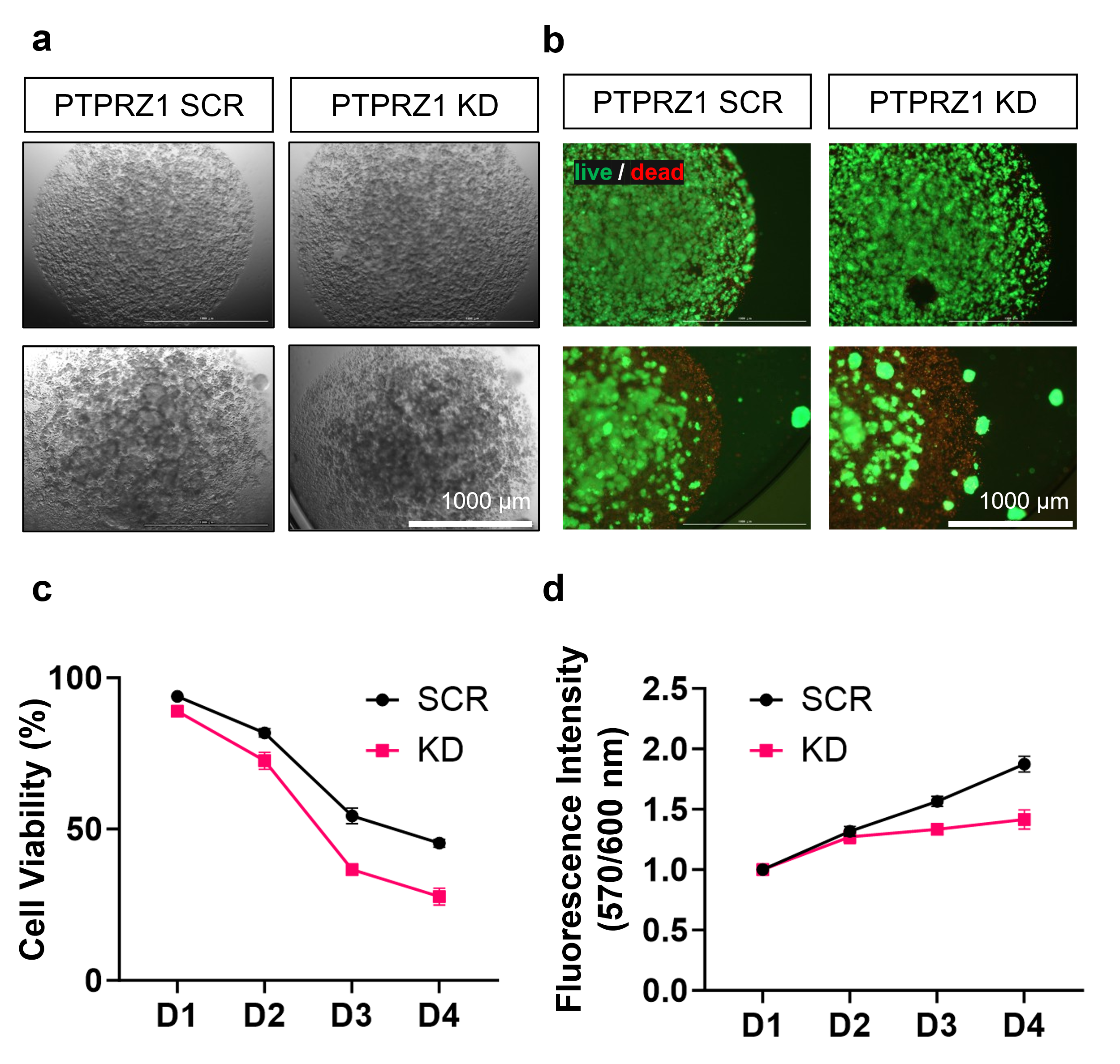

**Supplementary Fig. 8 | PTPRZ1 knockdown impairs hiPSC survival and metabolic activity under PTN-incorporated bioink conditions.**
**a,** Representative bright-field images of scrambled control (SCR) and PTPRZ1-knockdown (KD) hiPSC constructs cultured in PTN-incorporated bioink. Scale bar, 1000 μm. **b,** Representative Live/Dead fluorescence images of SCR and PTPRZ1-KD constructs, showing viable (green) and dead (red) cells. Scale bar, 1000 μm. **c,** Quantification of cell viability in SCR and PTPRZ1-KD constructs from days 1–4 after printing. **d,** Metabolic activity of SCR and PTPRZ1-KD constructs during 4 days of culture, assessed by fluorescence intensity at 570/600 nm. Data are presented as mean ± [SEM].

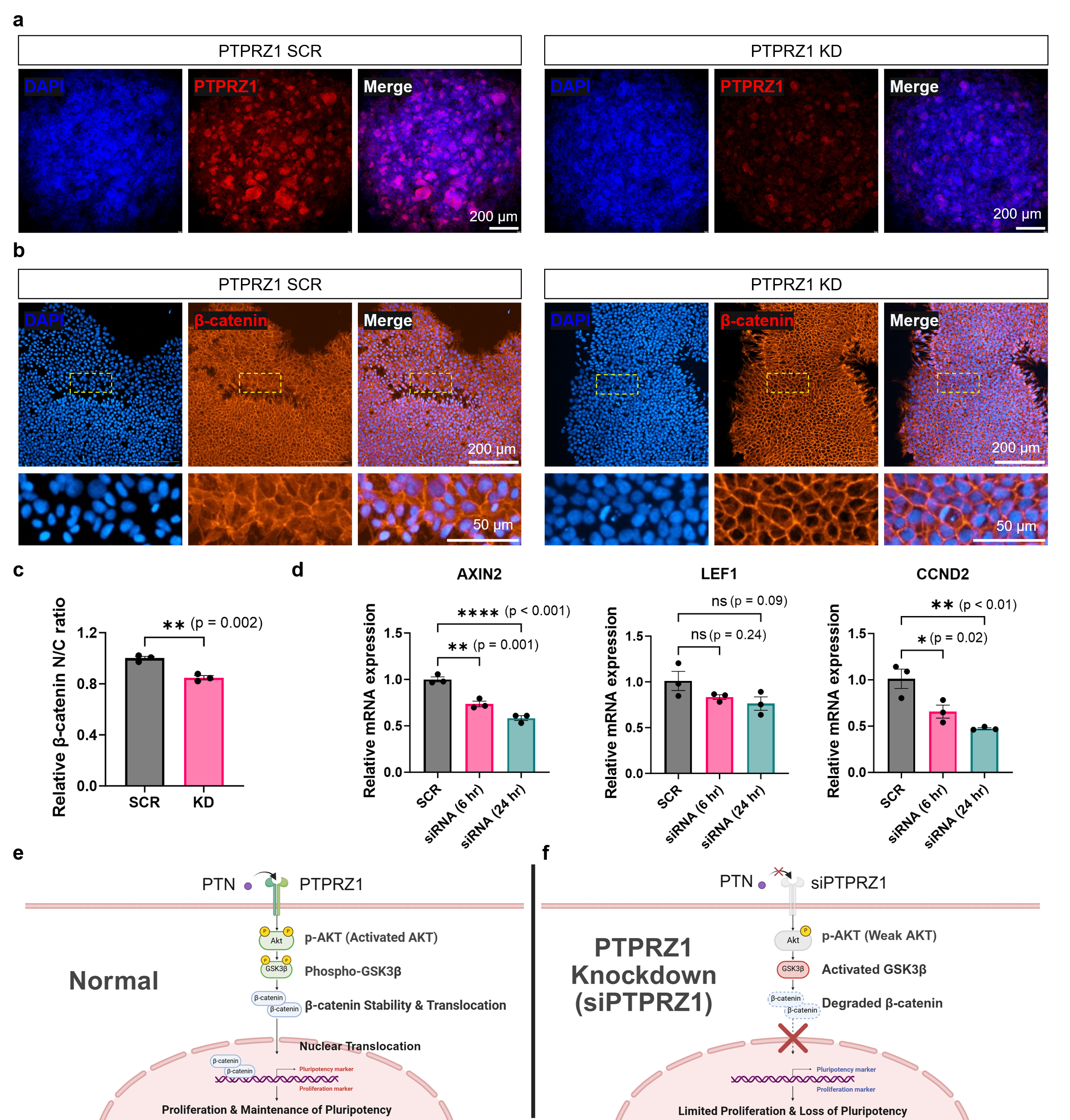

**Supplementary Fig. 9 | PTPRZ1 knockdown attenuates β-catenin nuclear localization and Wnt/β-catenin-associated transcriptional activity.**
**a,** Representative immunofluorescence images of PTPRZ1 expression in scrambled control (SCR) and PTPRZ1-knockdown (KD) hiPSC constructs. Nuclei were counterstained with DAPI. Scale bars, 200 μm. **b,** Representative immunofluorescence images of β-catenin localization in SCR and PTPRZ1-KD constructs. Boxed regions are shown at higher magnification in the lower panels. Nuclei were counterstained with DAPI. Scale bars, 200 μm (upper) and 50 μm (lower). **c,** Quantification of the nuclear-to-cytoplasmic (N/C) β-catenin intensity ratio in SCR and PTPRZ1-KD constructs. **d,** qRT–PCR analysis of the Wnt/β-catenin-associated genes *AXIN2*, *LEF1* and *CCND2* following PTPRZ1 knockdown at the indicated time points. Data are presented as mean ± SEM. Exact *P* values are shown in the graphs; ns, not significant. **e, f,** Proposed model of PTN–PTPRZ1-mediated signaling under control (**e**) and PTPRZ1-knockdown (**f**) conditions. PTN–PTPRZ1 signaling is proposed to promote AKT phosphorylation, GSK3β inhibition and β-catenin stabilization and nuclear translocation, thereby supporting proliferation and maintenance of pluripotency. PTPRZ1 knockdown is proposed to attenuate this signaling axis and reduce β-catenin activity.

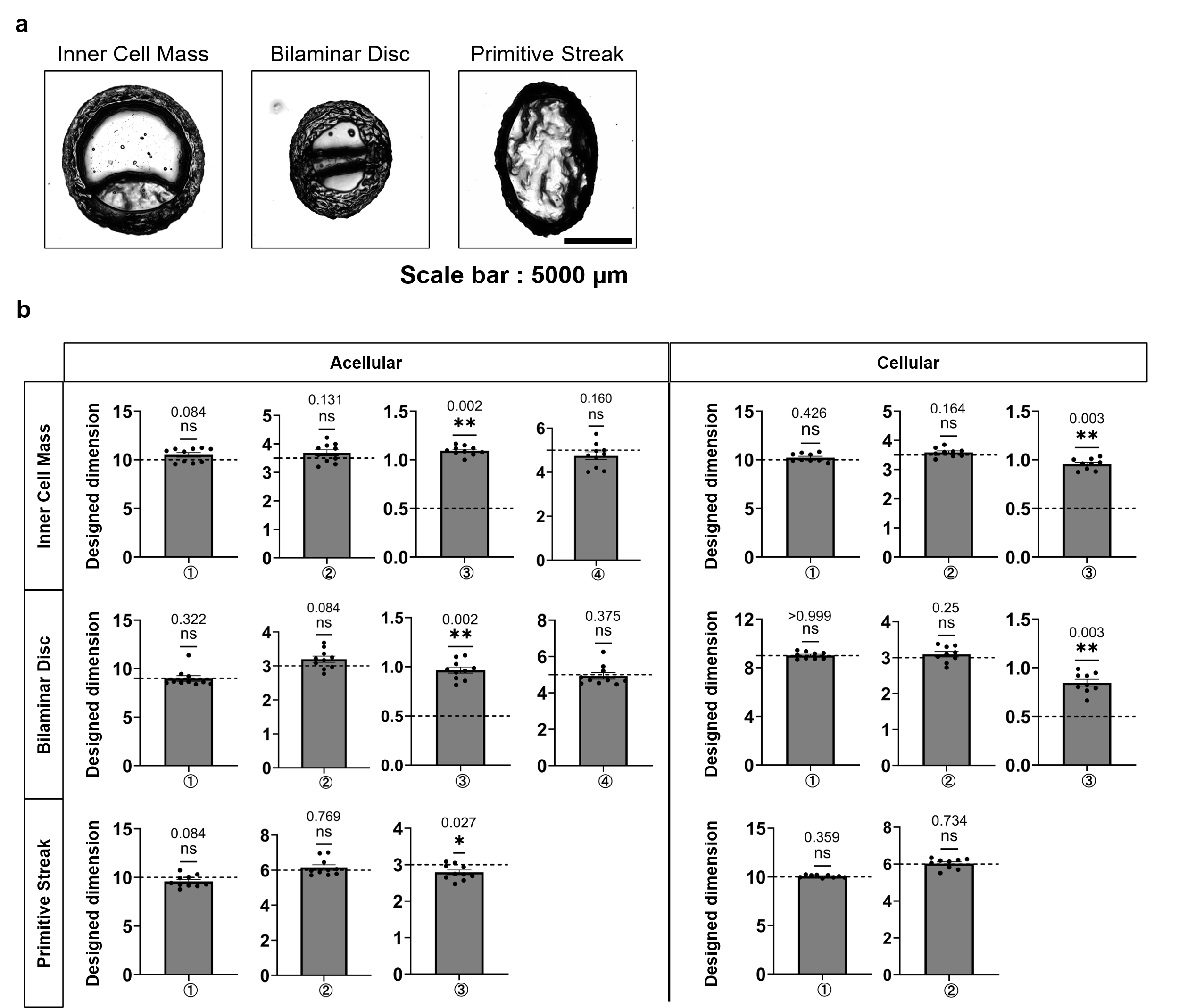

**Supplementary Fig. 10 | Dimensional fidelity of embryo-inspired constructs generated by 3D bioprinting.**
**a,** Representative images of acellular embryo-inspired constructs designed to mimic the inner cell mass (ICM), bilaminar disc and primitive streak geometries. Scale bar, 5000 μm. **b,** Quantitative comparison of the measured dimensions of acellular and cellular constructs with their corresponding designed dimensions. Measurements are shown for the indicated geometric parameters (①–④) of the ICM and bilaminar disc constructs and parameters ①–③ or ①–② of the primitive streak constructs, as applicable. Dashed horizontal lines indicate the predefined designed dimensions. Exact *P* values are shown above the graphs; ns, not significant; *P* < 0.05; **P* < 0.01.

**Supplementary Table 1 | KEGG pathway enrichment analysis of candidate ligands identified from epiblast-associated ligand–receptor interactions.**

**Pre-implantation**

| **KEGG_PATHWAY** | **Count** | **P-Value** | **Fold Enrichment** | **Receptor** | **Ligand** |
| --- | --- | --- | --- | --- | --- |
| Proteoglycans in cancer | 7 | 0.00000379 | 14.81 | ITGB1,SDC2,FZD8,ITGAV,ITGA5,CD44,FGFR1 | FN1, COL4A1, LAMA1, RSPO3, SPARC |
| ECM-receptor interaction | 5 | 0.0000382 | 24.25 | ITGB1,ITGA3,ITGAV,ITGA5,CD44 | FN1, COL4A1, LAMA1 |
| Arrhythmogenic right ventricular cardiomyopathy | 4 | 0.000848 | 20.08 | ITGB1,ITGA3,ITGAV,ITGA5 | FN1 |
| PI3K-Akt signaling pathway | 6 | 0.000962 | 7.15 | ITGB1,ITGA3,INSR,ITGAV,ITGA5,  FGFR1 | FN1, CALM2, SPARC |
| Hypertrophic cardiomyopathy | 4 | 0.00128 | 17.44 | ITGB1,ITGA3,ITGAV,ITGA5 | FN1 |
| Hematopoietic cell lineage | 4 | 0.00131 | 17.27 | ITGA3,CD9,ITGA5,CD44 | FN1, L1CAM, COL4A1, LAMA1 |
| Regulation of actin cytoskeleton | 5 | 0.0015 | 9.30 | ITGB1,ITGA3,ITGAV,ITGA5,FGFR1 | FN1, SPARC |
| Dilated cardiomyopathy | 4 | 0.00151 | 16.44 | ITGB1,ITGA3,ITGAV,ITGA5 | FN1 |
| Cytoskeleton in muscle cells | 5 | 0.00152 | 9.26 | ITGB1,ITGA3,SDC2,ITGAV,ITGA5 | FN1 |
| Signaling pathways regulating  pluripotency of stem cells | 4 | 0.00372 | 11.99 | FZD8,ACVR1B,ACVR2B,  FGFR1 | NODAL, TDGF1, RSPO3, SPARC |
| Virion - Hepatitis viruses | 3 | 0.00494 | 26.98 | CD81,CD9,LDLR | GPC3, L1CAM, HBEGF, APOA1 |
| Pathways in cancer | 6 | 0.00525 | 4.86 | ITGB1,ITGA3,FZD8,ITGAV,FGFR1,LRP6 | FN1, RSPO3, SPARC |
| Human papillomavirus infection | 5 | 0.00554 | 6.48 | ITGB1,ITGA3,FZD8,ITGAV,ITGA5 | FN1, RSPO3 |
| Wnt signaling pathway | 4 | 0.00631 | 9.92 | FZD8,GPC4,LGR6,LRP6 | RSPO3, COL18A1 |
| Focal adhesion | 4 | 0.00964 | 8.51 | ITGB1,ITGA3,ITGAV,ITGA5 | FN1 |
| Small cell lung cancer | 3 | 0.0177 | 13.92 | ITGB1,ITGA3,ITGAV | FN1 |
| Parathyroid hormone synthesis, secretion and action | 3 | 0.00554 | 6.48 | PTH1R,FGFR1,LRP6 | CGA, **SPARC**, RSPO3 |
| Fluid shear stress and atherosclerosis | 3 | 0.0387 | 9.12 | SDC2,ITGAV,ACVR2B | FN1, NODAL, TDGF1 |
| Breast cancer | 3 | 0.0418 | 8.75 | FZD8,FGFR1,LRP6 | RSPO3, SPARC |
| mTOR signaling pathway | 3 | 0.047 | 8.20 | INSR,FZD8,LRP6 | RSPO3, CALM2 |
| Phagosome | 3 | 0.0475 | 8.14 | ITGB1,ITGAV,ITGA5 | FN1 |
| Cell adhesion molecules | 3 | 0.0481 | 8.09 | ITGB1,SDC2,ITGAV | FN1 |
| Rap1 signaling pathway | 3 | 0.0789 | 6.11 | ITGB1,INSR,FGFR1 | FN1, CALM2, SPARC |
| Hormone signaling | 3 | 0.0835 | 5.91 | INSR,ITGAV,PTH1R | CALM2, FN1, CGA |

Post implantation

| **KEGG_PATHWAY** | **Count** | **P-Value** | **Fold Enrichment** | **Receptor** | **Ligand** |
| --- | --- | --- | --- | --- | --- |
| IgSF CAM signaling | 5 | 0.00216 | 8.27 | PTPRS,PTPRZ1,UNC5B,UNC5D,NECTIN2 | PTN, MDK, GNAI2, GPC3, NECTIN3 |
| Virion - Hepatitis viruses | 3 | 0.00364 | 31.24 | CD81,CD9,LDLR | GPC3, HBEGF, APOA1 |
| Signaling pathways regulating  pluripotency of stem cells | 3 | 0.0298 | 10.41 | FZD8,ACVR1B,ACVR2B | RSPO3, NODAL, TDGF1 |
| Cell adhesion molecules | 3 | 0.0362 | 9.37 | PTPRS,SDC2,NECTIN2 | PTN, FN1, APOE, NECTIN3 |
| Wnt signaling pathway | 3 | 0.0421 | 8.62 | FZD8,LGR6,LRP6 | RSPO3 |
| Proteoglycans in cancer | 3 | 0.0561 | 7.35 | SDC2,FZD8,CD44 | FN1, RSPO3, PTN, APOE, PKM |
| Malaria | 2 | 0.0907 | 19.99 | CD81,SDC2 | GPC3, FN1, PTN, APOE |
| Cholesterol metabolism | 2 | 0.0925 | 19.60 | ABCA1,LDLR | APOA1 |
